# The SPFH complex HflK-HflC regulates aerobic respiration in bacteria

**DOI:** 10.1101/2024.04.21.590321

**Authors:** Maria Isabel Perez-Lopez, Paul Lubrano, Georgia Angelidou, Timo Glatter, Nicole Paczia, Hannes Link, Victor Sourjik

## Abstract

The bacterial HflK-HflC membrane complex is a member of the highly conserved SPFH protein family, which is found throughout all domains of life and includes eukaryotic stomatins, flotillins, and prohibitins. These proteins organize cell membranes and are involved in various processes. However, the exact physiological functions of most bacterial SPFH proteins remain unclear. Here, we report that the HflK-HflC complex in *Escherichia coli* is required for growth under high aeration. The absence of this complex causes an aerobic growth defect due to a reduced abundance of IspG, a crucial enzyme in the isoprenoid biosynthetic pathway. This reduction leads to lower levels of ubiquinone, reduced respiration, lower ATP levels, and misregulated expression of respiratory genes. The regulation of aerobic respiration by the HflK-HflC complex resembles the mitochondrial respiratory defects caused by prohibitin mutations in mammalian and yeast cells, suggesting a functional commonality between these bacterial and eukaryotic SPFH proteins.

## INTRODUCTION

Members of the SPFH (<u>S</u>tomatin, <u>P</u>rohibitin, <u>F</u>lotillins, and <u>H</u>flK-HflC) protein family have been identified in all three domains of life^1, 2^. A common feature of these membrane proteins is an evolutionarily conserved prohibitin homology (PHB) domain (also called SPFH domain), which may have lipid-protein binding properties^3^. The SPFH proteins share a common property of self-oligomerization into large membrane-spanning or membrane-anchored complexes, and they appear to have diverse but poorly understood functions, mostly related to the organization of lipid membranes^4–6^.

In eukaryotic cells, SPFH proteins are present at various cellular locations, including the plasma membrane, Golgi apparatus, mitochondria, and endoplasmic reticulum^3, 7^, where they play an important role in scaffolding proteins and specific lipids within lipid domains. The SPFH proteins are involved in various biological processes, with stomatins contributing to the regulation of ion channels^8, 9^, and flotillins being associated with signal transduction, endocytosis, and neuronal regeneration^7, 10, 11^. Prohibitins, located in the inner mitochondrial membrane, form large hetero-oligomers that interact with the AAA+ membrane protease^12^. The absence of prohibitins affects several cellular processes, including cell proliferation, apoptosis, and respiration, but the mechanisms behind these effects are still unknown^13–16^.

Bacterial SPFH family proteins were described more than two decades ago^1^, but their functions are even less understood than those of their eukaryotic counterparts. Research on Gram-positive bacteria has revealed certain structural and functional similarities between eukaryotic and bacterial flotillins^17^, with the scaffolding activity of these bacterial flotillins being important for the regulation of membrane fluidity and the assembly of protein complexes involved in signal transduction^18–20^. Even less is known about the functions of SPFH proteins in Gram-negative bacteria. In *Escherichia coli*, four proteins containing the PHB domain have been identified: QmcA, YqiK, and the complex HflK-HflC (=HflKC), all of which are localized in the inner membrane. While the functions of QmcA and YqiK remain unclear, the HflKC complex is known to interact with FtsH, an integral membrane ATP-dependent Zn^2+^ metalloprotease belonging to the AAA+ family of ATPases^21^. HflK and HflC have a similar secondary structure consisting of a single transmembrane helix at the N-terminus followed by large periplasmic SPFH1 and SPFH2 domains and coiled-coil domains. HflK, HflC, and FtsH form a large structure consisting of 12 copies of the HflKC heterodimer, providing a large compartmentalized cage for four embedded FtsH hexamers^22, 23^. This complex shares features with the multimeric assemblies formed by eukaryotic prohibitins in the inner membrane of mitochondria interacting with a hexameric AAA+ protease homologous to FtsH^24, 25^.

FtsH degrades membrane and cytoplasmic proteins involved in several cellular pathways^26, 27, 28^, and deletion of the *ftsH* gene causes a severe growth defect^29^. In contrast, no pronounced growth phenotype has been reported for *E. coli* lacking the HflKC complex^30^, and the physiological significance of this complex, including the HflKC-dependent regulation of FtsH, remains unclear^28^. Here, we demonstrate that the HflKC complex is necessary for growth under conditions of high aeration. This effect could be explained by a decrease in the abundance of IspG, a key enzyme in the isoprenoid biosynthesis pathway, resulting in reduced levels of ubiquinone, which is essential for aerobic respiration. These findings reveal a novel function of the HflKC complex in aerobic respiration, which may be analogous to the function of eukaryotic prohibitins in mitochondria.

## RESULTS

### HflKC complex is important for *E. coli* growth under high aeration

When an *E. coli* strain deleted for the *hfl* genes was phenotyped under various conditions, it exhibited a growth defect that was dependent on aeration and medium composition. When *E. coli* was cultured in rich tryptone broth (TB) medium on an orbital shaker, the growth of both single and double deletions of the *hflK* and *hflC* genes was similar to that of the wild-type strain at low shaking rates (Figures 1A and S1A). However, at higher shaking rates, the growth of the *ΔhflK ΔhflC* (= *ΔhflK*C) strain was significantly slower than that of the wild-type strain (Figures 1B-1D and S1B, S1C). While wild-type growth increased at higher shaking rates, as expected from better aeration, growth of the *ΔhflKC* mutant actually decreased. A weaker but similar growth defect was observed for the *ΔhflK* strain, whereas the *ΔhflC* strain did not differ from wild-type growth. The observed growth defect of the *ΔhflKC* strain was specific, as it could be largely complemented by co-expressing the *hflK* and *hflC* genes from a plasmid (Figures 1E and 1F).

**Figure 1.**
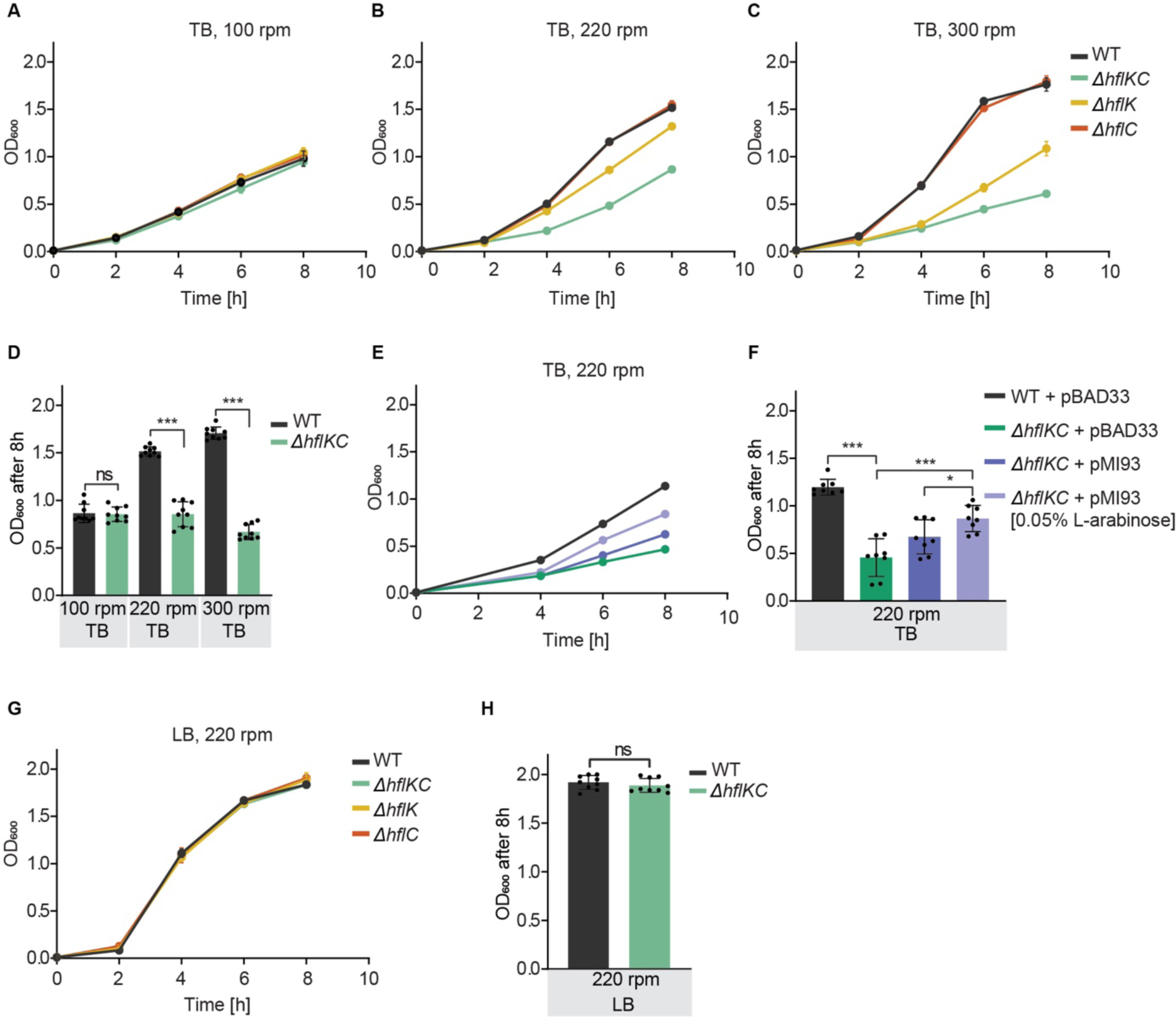
HflKC complex is important for *E. coli* growth under high aeration. (A-D) Growth of *E. coli ΔhflKC, ΔhflK*, and *ΔhflC* strains and corresponding wild-type (WT) in TB medium at 100 rpm (A), 220 rpm (B), or 300 rpm (C) shaking rate, quantified by optical density at 600 nm (OD_600_), and final OD_600_ after 8 h of growth (D). (E, F) Growth of *ΔhflKC* and WT strains carrying either an empty vector (pBAD33) or the pBAD33-derived expression plasmid pMI93 encoding *hflK* and *hflC*, in TB at 220 rpm (E) and corresponding final OD_600_ (F). Where indicated, 0.05% L-arabinose was added to induce expression. (G, H) Growth of *E. coli ΔhflK, ΔhflC, ΔhflKC,* and WT strains in LB at 220 rpm (G) and corresponding final OD_600_ (H). For these and other growth curves, data represent the mean and standard deviation (SD) of three independent cultures grown in the same representative experiment. See Figure S1A-S1D for additional biological replicates. For final OD_600_ comparisons, data represent the mean and SD of independent cultures, indicated by dots, grown in three different experiments. Significance of indicated differences between samples: *p<0.05, ***p<0.001, and ns = not significant by unpaired *t*-test.

These results indicate that the absence of the HflKC complex or of HflK causes a specific aeration-dependent growth phenotype. Interestingly, however, no growth defect was observed for the *ΔhflK* and *ΔhflKC* strains at high aeration in an even richer Luria-Bertani (LB) medium (Figures 1G, 1H and S1D), which contains yeast extract in addition to the tryptone and NaCl that are present in both LB and TB. We therefore tested whether the addition of a fermentable carbon source to TB could restore the growth of the *ΔhflKC* mutant. However, while supplementation of TB with glucose resulted in faster growth, the difference between the *ΔhflKC* strain and the wild type remained (Figures S1E and S1F). The growth phenotype of the *ΔhflKC* strain further remained evident when cells were cultured at high aeration in M9 minimal medium containing glucose as the sole carbon source (Figures S1G and S1H). Consistent with the aeration dependence of the growth defect observed for the *ΔhflKC* strain, no difference in growth from the wild type was observed in TB under anaerobic conditions (Figures S1I and S1J).

### Absence of HflKC complex affects the abundance of respiration-related proteins

To identify possible causes for the observed growth defect, we first analyzed changes in whole-cell protein levels caused by deletion of the *hflK* and *hflC* genes for *E. coli* cultures grown in LB or TB under strong shaking. Consistent with similar growth of the Δ*hflKC* and wild-type strains in LB (Figure 1G), only a small number of proteins showed pronounced differences in abundance under these conditions (Figure 2A; Tables 1 and S1). In contrast, differences between cultures grown in TB, where the deletion strain showed a growth defect at high aeration (Figure 1B), were much more extensive (Figure 2B; Table S2). Fewer differences in protein composition were observed when the two strains were grown under anaerobic conditions (Figure 2C; Table S3), consistent with their similar growth (Figure S1I).

**Figure 2.**
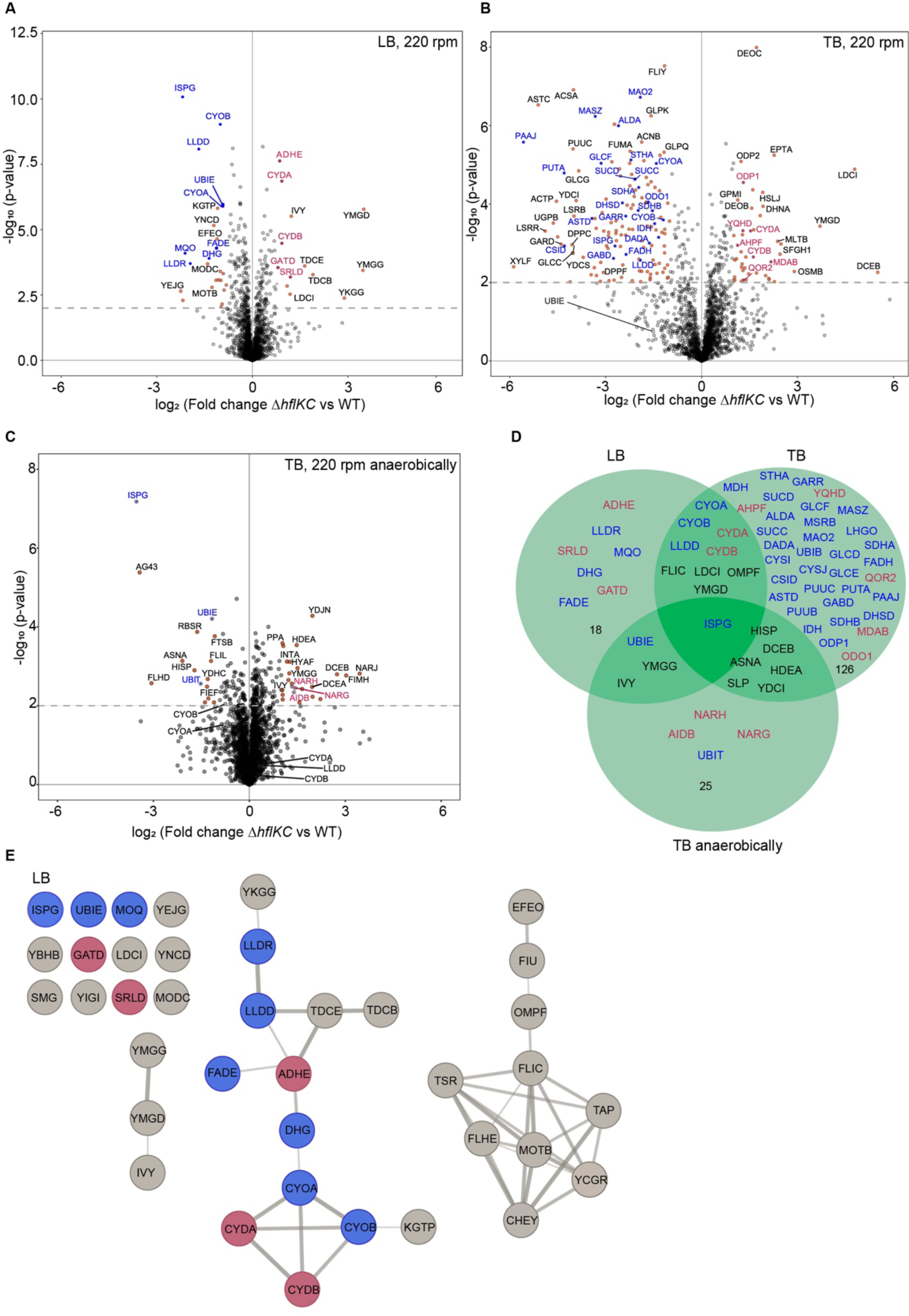
Absence of HflKC complex affects the abundance of respiration-related and other proteins. (A-C) Difference in protein levels between *ΔhflKC* and WT strains. Cultures were grown in LB (A), TB (B), or anaerobically in TB (C). Data represent six (LB) or three (TB) independent cultures. Proteins with differences in expression that were considered significant (see also Tables 1, S1, S2, S3) are labeled, with respiration-related proteins highlighted in either blue (downregulated) or red (upregulated). (D) Commonalities and differences between proteins significantly up- or downregulated in *ΔhflKC* under different conditions. Colors of protein labels are the same as in other panels. Respiration-related proteins and those affected under more than one condition are shown, and the number of other proteins affected under a particular condition is shown. (E) The STRING diagram showing proteins that are significantly up- or downregulated in the *ΔhflKC* deletion strain, with links indicating relationships between proteins. Proteins related to respiration are colored in red (upregulated) or blue (downregulated).

**Table 1.** Respiratory proteins showing significant differences between *ΔhflKC, ΔhflK,* or *ΔhflC* and wild-type strains during growth in LB at 220 rpm.

| Protein | Function | Log<sub>2</sub> (Fold change) |  |  |
| --- | --- | --- | --- | --- |
| | | $\Delta hflKC$ vs<br>WT | $\Delta hflK$ vs<br>WT | $\Delta hflC$ vs<br>WT |
| ISPG | Oxidoreductase involved in isoprenoid biosynthesis | -2.28 | -2.37 | 0.02 |
| MQO | Malate: quinone oxidoreductase | -2.15 | -1.84 | -0.33 |
| LLDR | L-lactate dehydrogenase operon regulator | -1.90 | -2.26 | -0.44 |
| LLDD | L-lactate dehydrogenase | -1.66 | -1.52 | -0.38 |
| DHG | Quinoprotein glucose dehydrogenase | -1.34 | -1.26 | -0.23 |
| FADE | Acyl-CoA dehydrogenase | -1.13 | -0.50 | -0.78 |
| CYOB | Cytochrome bo(3) ubiquinol oxidase subunit 1 | -1.01 | -1.01 | -0.20 |
| UBIE | Ubiquinone biosynthesis | -0.92 | -0.93 | -0.01 |
| CYOA | Cytochrome bo(3) ubiquinol oxidase subunit 2 | -0.91 | -0.85 | 0.00 |
| SRLD | Sorbitol-6-phosphate 2-dehydrogenase | 1.18 | 1.44 | -0.36 |
| CYDA | Cytochrome bd-I ubiquinol oxidase subunit 1 | 0.96 | 0.85 | -0.03 |
| ADHE | Fused acetaldehyde-CoA dehydrogenase | 0.94 | 0.89 | -0.08 |
| CYDB | Cytochrome bd-I ubiquinol oxidase subunit 2 | 0.91 | 0.68 | -0.05 |
| GATD | Galactitol-1-phosphate 5-dehydrogenase | 0.82 | 1.12 | 0.28 |
| DMSA | Dimethyl sulfoxide reductase subunit A | 0.26 | -0.95 | 0.40 |

Data from Figures 2A and S3A, S3B. Differences with p-value <0.05 and log2 of fold change >0.8 were considered significant.

Despite this dependence on incubation conditions, the levels of several proteins showed consistent differences between the Δ*hflKC* and wild-type strains (Figure 2D). Among the proteins whose abundance was significantly perturbed under aerobic conditions in both LB and TB were two cytochrome quinol oxidases, CyoABCD (*bo_3_*) and CydAB (*bd*), which are used by *E. coli* under aerobic (i.e., high O_2_) and microaerobic (low O_2_) conditions, respectively^31^. The levels of two cytochrome quinol oxidases showed opposite changes, with the catalytic subunits CyoAB of the aerobic quinol oxidase *bo_3_* being reduced in the Δ*hflKC* strain, whereas the levels of the microaerobic quinol oxidase CydAB were increased. The expression of several other respiration-related proteins was also affected in LB (Figure 2E; Table S1), and even more so in TB under aerobic conditions (Table S2).

We also observed a strong reduction in the levels of two metabolic enzymes, UbiE and IspG, which are involved in the biosynthesis of respiratory chain electron carriers. UbiE methyltransferase is part of the ubiquinone and menaquinone biosynthetic pathway^32^. IspG belongs to the methylerythritol phosphate (MEP) pathway and catalyzes the conversion of ME-cPP (2C-methyl-D-erythritol 2,4-cyclodiphosphate) to HMBPP (hydroxymethylbutenyl 4-diphosphate), a key substrate for the production of isoprenoids, which are also required for quinone biosynthesis^33^ (Figure 3A). The reduced abundance of these two enzymes was also observed even under anaerobic conditions and thus independent of the respiratory status of the *E. coli* cells. Notably, although the change in UbiE level was below the significance threshold in TB under aerobic conditions, its expression was still reduced (Figure 2B).

**Figure 3.**
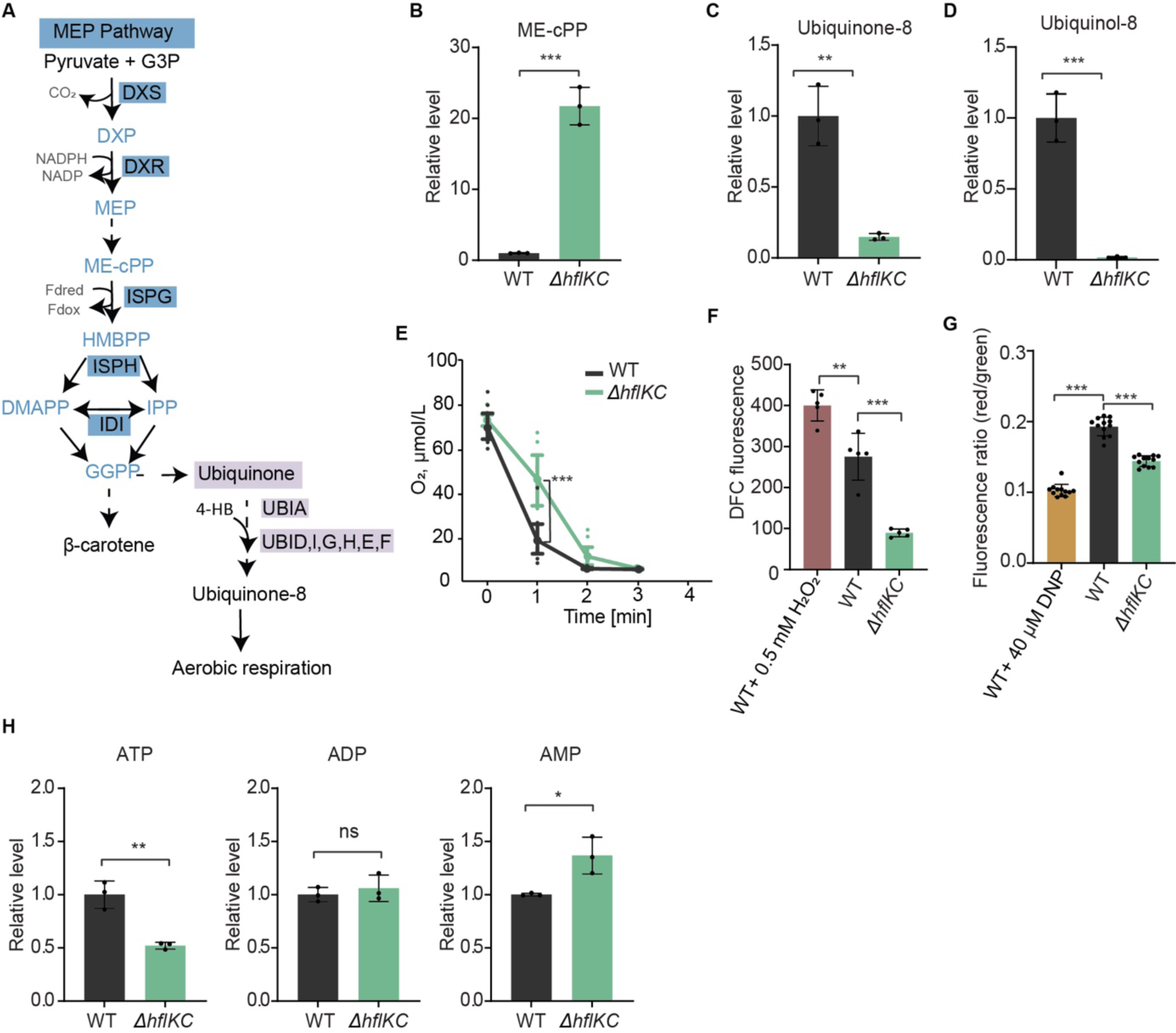
*ΔhflKC* strain shows reduced ubiquinone levels, aerobic respiration, and ATP levels. (A) Methyl-D-erythritol phosphate (MEP) pathway in *E. coli.* Metabolic intermediates are colored in light blue, and selected enzymes are shown on either dark blue (MEP pathway) or purple (ubiquinone biosynthesis) background. (B-D) Levels of the IspG substrate ME-cPP (B) and of ubiquinone-8 (C) and ubiquinol-8 (D) in *ΔhflKC* relative to the WT strain. Strains grown at 220 rpm in either M9 glucose minimal medium (B) or in TB (C, D). Data represent the mean and SD of three independent cultures. (E) Oxygen consumption by WT and *ΔhflKC* cells. Cultures were grown in TB at 220 rpm, resuspended in fresh TB, and changes in the levels of dissolved oxygen were quantified over time. Large symbols represent the mean and SD of eight independent measurements (shown by small dots) for cells from one culture. See also Figure S4. (F) Levels of ROS in WT and *ΔhflKC* cells grown in TB at 220 rpm, measured using the DCF fluorescent probe as illustrated in Figure S5. Treatment with hydrogen peroxide (H_2_O_2_) was used as a positive control for elevated ROS levels. Data represent the mean and SD of five measurements with 30,000 cells per measurement. (G) Membrane potential of WT and *ΔhflKC* cells grown in TB at 220 rpm, measured using the DiOC_2_(3) dye as illustrated in Figure S6. Dinitrophenol (DNP) was used as a control. Data represent the mean and SD of twelve measurements from two independent experiments with 30,000 cells per measurement. (H) Levels of ATP, ADP, and AMP (H) in cells grown in M9 glucose minimal medium at 220 rpm. Means of three independent cultures and SD are shown. Significance of indicated differences between samples: * p<0.05, ** p<0.01, *** p<0.001, and ns = not significant by unpaired *t*-test.

In addition to the cluster of respiration-related proteins, significant changes in the levels of other proteins were also observed in the Δ*hflKC* strain. In particular, proteins involved in motility and chemotaxis were downregulated in LB (Figure 2E; Table S1) and also in TB under both aerobic and anaerobic conditions (Tables S2 and S3). Notably, the abundance of known FtsH substrates^34^ and of FtsH itself was not significantly affected in either LB or TB (Figures S2A and S2B), confirming that the Δ*hflKC* deletion does not lead to a general change in FtsH activity.

Although our primary focus was on the phenotype of the strain lacking the entire HflKC complex, we also evaluated the individual effects of the *hflK* and *hflC* deletions. Consistent with their growth, the proteome profiles of the Δ*hflKC* and Δ*hflK* strains were similar (Figures 2A, 2B and S3A, S3C), whereas the Δ*hflC* strain showed little change in proteome composition compared to the wild type (Figures S3B and S3D). Thus, the phenotype observed in the Δ*hflKC* strains appears to be primarily due to the absence of HflK, whereas the absence of HflC can be tolerated by the cell and becomes apparent only in the background of the *hflK* deletion. Notably, both single deletions of *hflK* and *hflC* caused a reduction in the level of the other component of the HflKC complex (Figures S3A-S3D), but such a reduction in the case of the Δ*hflC* strain was apparently not sufficient to cause the growth phenotype or to affect proteome composition.

### *ΔhflKC* strain shows reduced ubiquinone levels, aerobic respiration, and ATP levels

Given the greatly reduced levels of IspG in the *ΔhflKC* strain and the importance of the MEP pathway for the ubiquinone biosynthesis (Figure 3A), we examined the impact of the *ΔhflKC* deletion on the MEP pathway and on ubiquinone levels. Consistent with low IspG activity, the level of the IspG substrate ME-cPP was largely elevated in the *ΔhflKC* strain compared to the wild type (Figure 3B), whereas the levels of the oxidized (ubiquinone-8) and especially the reduced (ubiquinol-8) forms of ubiquinone were strongly reduced (Figures 3C and 3D). Thus, the downregulation of IspG, and possibly also of UbiE downstream in the pathway (Figure 3A), apparently causes a disruption in the ubiquinone biosynthesis in the absence of the HflKC complex.

Because low levels of ubiquinone could cause a reduction in aerobic respiratory activity, we compared the consumption of dissolved oxygen by the *ΔhflKC* and wild-type cell cultures. Indeed, oxygen consumption by the *ΔhflKC* cell culture was significantly reduced (Figures 3E and S4). Further consistent with reduced respiration, the level of reactive oxygen species (ROS) assessed using the dichlorodihydrofluorescein (DCF) probe (Figure 3F and S5), as well as the membrane potential assessed using the 3,3’-diethyloxacarbocyanine iodide DiOC_2_(3) probe (Figure 3G and S6) were also reduced in *ΔhflKC* cells.

Such reduced respiration and the resulting decrease in membrane potential could lead to reduced ATP production in *ΔhflKC* cells. This decrease was indeed evident when the levels of ATP, ADP, and AMP were quantified in *ΔhflKC* and wild-type cultures using targeted metabolomics. We observed that the level of ATP was lower and the level of AMP was higher in *ΔhflKC* cells, whereas the level of ADP remained unchanged (Figure 3H). These changes in the levels of adenosine phosphate groups could result in a lower energy charge in *ΔhflKC* cells.

### Reduced levels of IspG account for the respiratory phenotype of the *ΔhflKC* strain

Collectively, our data suggest that the lower ubiquinone levels, and consequently reduced aerobic respiration and poor growth at high aeration, may be due to low levels of IspG and/or UbiE. Since the reduction in IspG abundance was more pronounced and consistent across data sets, we hypothesized that it might be the primary cause of the observed respiratory phenotype. Indeed, induced expression of IspG from a plasmid restored ubiquinone (Figure 4A) and ubiquinol (Figure 4B) levels in *ΔhflKC* cells, as well as their oxygen consumption (Figures 4C and S7A), to wild-type strain levels. Growth of the *ΔhflKC* strain at high aeration (Figures 4D and 4E) and cell membrane potential (Figure 4F) also increased upon induction of IspG expression, even exceeding the wild type levels. Thus, all the observed respiration-related phenotypes of the *ΔhflKC* strain could be complemented by the overexpression of IspG.

**Figure 4.**
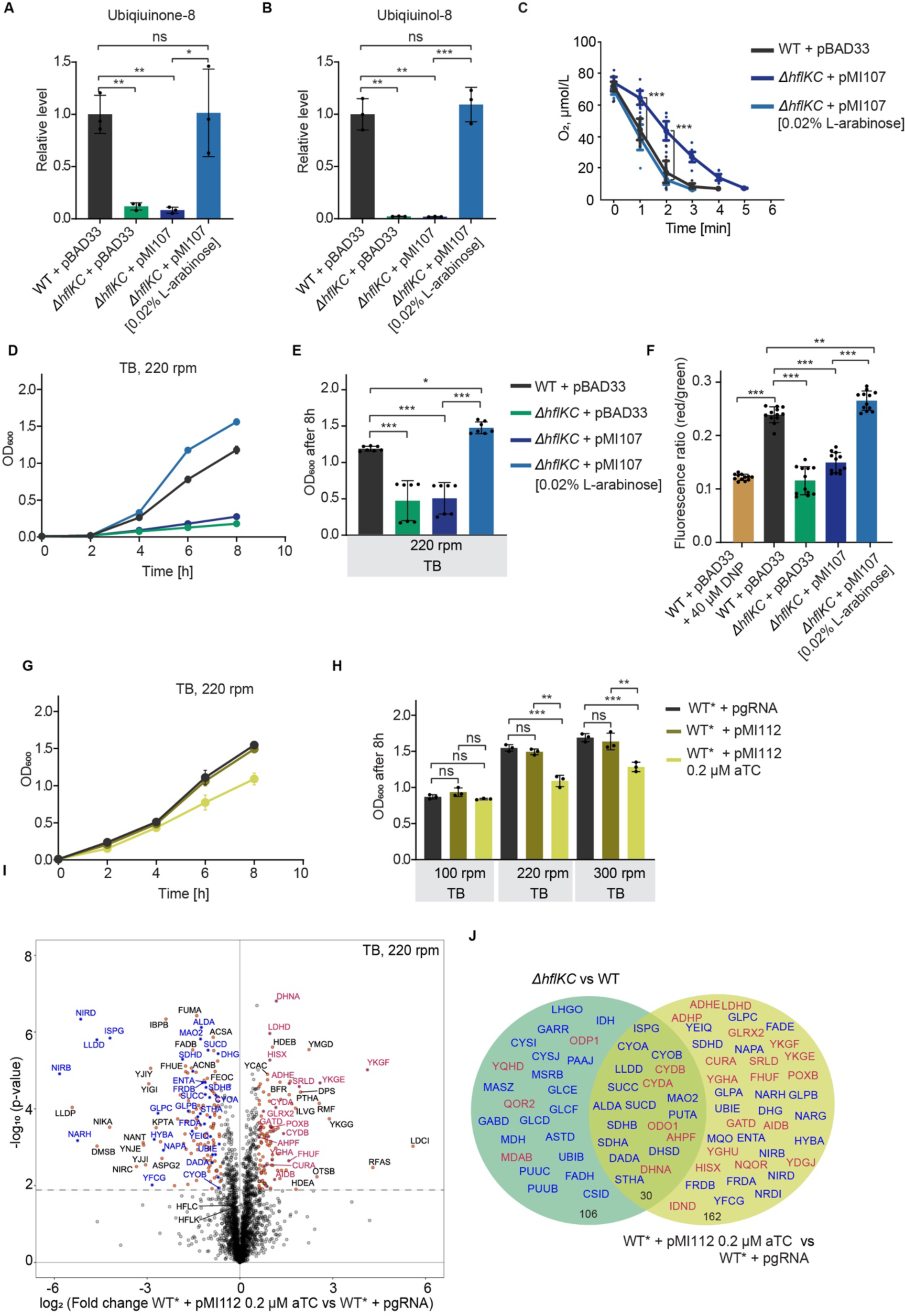
Reduced IspG levels cause the respiratory phenotype of the *ΔhflKC* strain. (A, B) Levels of ubiquinone-8 (A) and ubiquinol-8 (B) in the *ΔhflKC* strain, expressing IspG from an inducible plasmid vector, relative to WT strain carrying pBAD33. WT or *ΔhflKC* strains, transformed with empty vector pBAD33 or with pMI107 encoding *ispG* were grown in TB at 220 rpm; 0.02% L-arabinose was added to induce expression where indicated. Data represent the mean and SD of three independent cultures. (C) Oxygen consumption by the indicated strains. Measurements were performed as in Figure 3E. Large symbols represent the mean and SD of eight independent measurements for cells from one culture. See also Figure S7A. (D, E) Growth of the indicated strains (D) and corresponding final OD_600_ (E). (F) Measurements of membrane potential in the indicated strains, performed using the DiOC_2_(3) dye as in Figure 3G. (G, H) Growth of *E. coli* YYdCas9 (WT*) strain carrying the empty pgRNA vector or the pgRNA-derived pMI112 construct expressing guide RNA for *ispG* knockdown under the constitutive promoter (G) and final OD_600_ at indicated shaking rates (H). dCas9 expression was induced with 0.02 µM aTC (anhydrotetracycline). (I) Difference in protein levels between WT* carrying either pMI112 or pgRNA vector. Data are from three independent cultures. Proteins whose levels were considered to be significantly different between the two strains are labeled as in Figure 2. See also Table S3. (J) Commonalities and differences between proteins that are significantly up- or downregulated during growth in TB upon *hlfKC* deletion (Figure 2B) and upon *ispG* knockdown. Labels are as in Figure 2D. Significance of indicated differences between samples: * p<0.05, ** p<0.01, *** p<0.001, and ns = not significant by unpaired *t*-test.

Because *ispG* is essential in *E. coli*, we used dCas9 *ispG* knockdown to assess the effect of reduced IspG levels. This knockdown had no effect on *E. coli* growth at low aeration (Figure S7B), but reduced growth at high aeration (Figures 4G, 4H and S7C), effectively phenocopying the effects of *ΔhflKC* deletion. Changes in the abundance of several respiration-related proteins due to *ispG* knockdown were similar to those in the *ΔhflKC* strain (Figures 4I, 4J; Table S4), including reduced levels of UbiE and CyoAB and increased levels of CydAB. In contrast, the levels of motility-related and some other proteins were not affected by *ispG* knockdown, suggesting that their changes are unrelated to the reduced IspG levels.

### Changes in the abundance of respiratory proteins are caused by activation of the ArcAB system

Finally, we investigated the mechanism responsible for the observed global changes in the abundance of respiratory proteins due to reduced levels of IspG. In *E. coli*, the levels of (oxidized) quinones are known to repress the two-component ArcAB system^35^. The latter, in turn, controls the expression of a large number of respiration-related genes to mediate the transition from aerobic to anaerobic growth^36, 37^. Thus, we hypothesized that the reduced ubiquinone biosynthesis in the *ΔhflKC* might cause activation of the ArcAB system, leading to downregulation of aerobic respiratory genes and induction of the microaerobic cytochrome oxidase *bd-*I.

Indeed, although deletion of the *arcB* gene, which encodes the sensory kinase, itself negatively affected growth, we observed no additional impact of deletion of the *hflKC* genes in the *ΔarcB* background on aerobic growth in TB (Figures 5A, 5B and S8A, S8B). Furthermore, the changes in proteome composition caused by *arcB* deletion were largely opposite to those caused by *hflKC* deletion (Figures 5C and 5D; Table S5), and no changes in the levels of CyoAB or CydAB proteins were observed when comparing *ΔarcB* and *ΔhflKC ΔarcB* strains (Figures 5E and F; Table S6). This is consistent with our hypothesis that observed changes in the levels of respiratory proteins are dependent on the ArcAB system (Figure 5G). In contrast, the downregulation of IspG and UbiE, as well as of several other proteins, including those involved in motility, appears to be independent of the ArcAB system.

**Figure 5.**
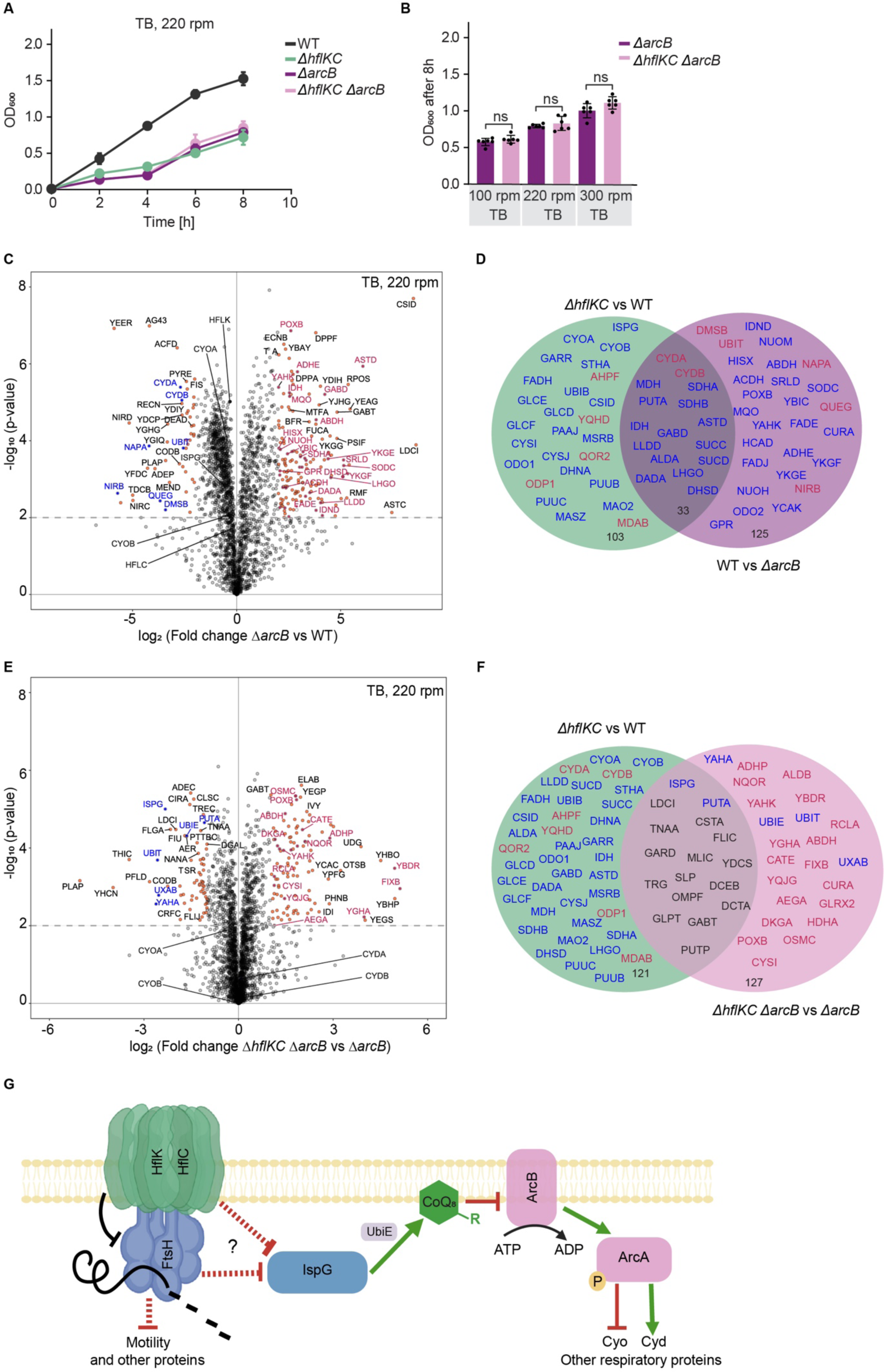
Changes in the abundance of respiratory proteins are caused by activation of the ArcAB system. (A, B) Growth of the WT, *ΔhflKC, ΔarcB and ΔhflKC ΔarcB* strains (A) and final OD_600_ at indicated shaking rates (B). ns = not significant by unpaired *t*-test. (C) Difference in protein levels between *ΔarcB* and WT strains. Data are for three independent cultures. Proteins whose levels were considered to be significantly different between the two strains are labeled as in Figure 2. See also Table S4. (D) Commonalities and differences between proteins that are significantly up- or downregulated during growth in TB upon *hflKC* deletion (Figure 2B) and upon *arcB* deletion (C). Labels are as in Figure 2D, but the sign of changes upon *arcB* deletion is inverted. (E) Difference in protein levels between the *ΔhflKC ΔarcB* and *ΔarcB* strains strains. Data are for three independent cultures. Labels are as in Figure 2. See also Table S5. (F) Commonalities and differences between proteins that are significantly up- or downregulated during growth in TB at 220 rpm upon *hflKC* deletion or in *ΔarcB* strain. Labels are as in Figure 2D. (G) Schematic representation of proposed function of the HflKC complex in respiration control. See text for details.

### The decrease in IspG levels is partly explained by its lower stability in the *ΔhflKC* strain

To better understand the possible origin of the reduced abundance of IspG in the *ΔhflKC* strain, we first compared *ispG* transcript levels between the *ΔhflKC* and wild-type strains grown in LB at 220 rpm. Our RT-PCR analysis revealed no significant difference (Figure 6A; Table S7), ruling out transcriptional regulation as a cause of the reduced IspG levels. Since the HflKC complex interacts with FtsH, an alternative explanation could be an increased degradation of IspG in the absence of this complex. We therefore examined the stability of IspG in both the wild-type and *ΔhflKC* strains. This was done by quantifying changes in the levels of IspG in bacterial cultures incubated in LB for varying periods of time in the presence of the translation inhibitor chloramphenicol. While no reduction was observed in the wild-type cells, suggesting that IspG is stable in the presence of the HflKC complex, a significant decrease in IspG abundance was observed after 30 min and 60 min in the *ΔhflKC* strain (Figures 6B and S9A). Thus, in the absence of the HflKC complex, IspG degradation is moderately but significantly increased. A similar decrease in IspG stability in the *ΔhflKC* strain compared to the wild type was observed in cultures grown in TB at 100 rpm (Figures S9B and S9C).

**Figure 6.**
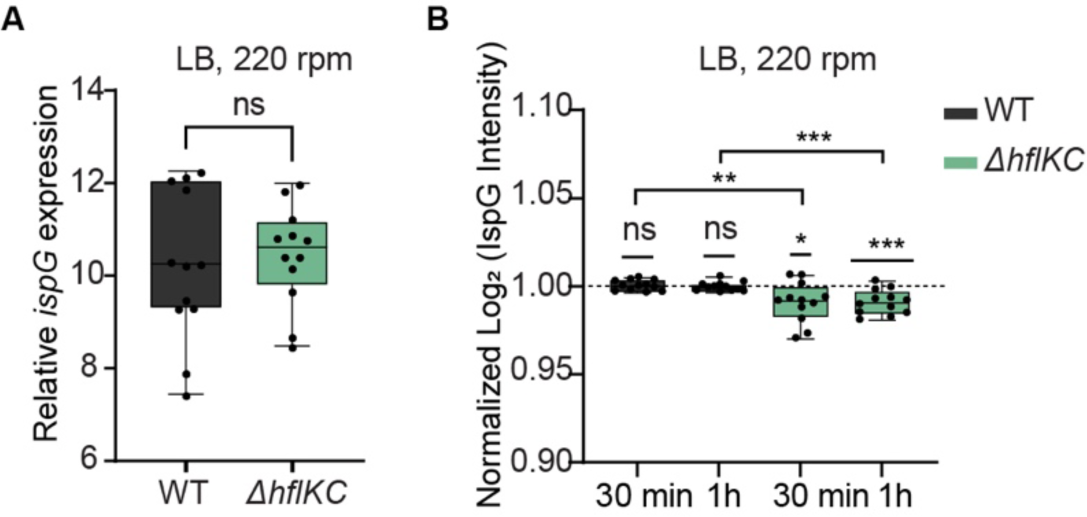
Decreased stability of IspG in the *ΔhflKC* strain. (A) Quantification of *ispG* transcript level in *ΔhflKC* and WT strains using real-time PCR (RT-PCR). The relative mRNA level of *ispG* is quantified as the Cq value and normalized to the Cq value for the housekeeping gene *ssrA*. Data represent the mean and SD for three independent RNA samples with quadruplicate measurements each. See also Table S7. (B) Changes in abundance of IspG in the WT and *ΔhflKC* strains upon incubation with chloramphenicol in LB at 220 rpm for 30 minutes or 1 hour. Abundance of IspG was determined by proteomics, and normalized to the initial time point. Data represent the mean and SD of two independent experiments with six independent cultures each. See also Figure S9A. Significance of the difference from the value of 1 by one sample *t*-test is indicated, as well as the significance of the difference between WT and *ΔhflKC* values at the same time point (indicated by brackets): *p< 0.05, **p< 0.01, ***p< 0.001, and ns = not significant.

## DISCUSSION

Although SPFH proteins are conserved between prokaryotes and eukaryotes, suggesting their fundamental importance for cellular function, the specific roles of these proteins remain poorly understood^2, 38^. In particular, only a few examples of the functional importance of SPFH proteins have been reported in prokaryotes^39–42^. Studies of SPFH proteins in *E. coli* have so far identified mild phenotypes that have not been mechanistically explained^30, 43^. This is particularly surprising for the HflKC complex, which is known to form a large oligomeric inner membrane cage that encloses the nearly essential AAA-type protease FtsH^22, 23^ and is thought to regulate FtsH access to its substrates^23^.

Here we demonstrate that the HflKC complex plays an important role during the growth of *E. coli* under conditions of high aeration. Our results suggest that the growth defect of the *ΔhflKC* strain under high aeration could be largely explained by a reduction in the level of IspG, an enzyme in the methylerythritol phosphate (MEP) pathway for isoprenoid biosynthesis (Figure 5G). The MEP pathway provides essential precursors for several cellular processes^44^, including the biosynthesis of pigments and ubiquinone^45–47^. Indeed, the level of ubiquinone-8 was greatly reduced in *ΔhflKC* cells. In addition to limiting the precursor supply for ubiquinone biosynthesis, the low level of IspG may further decrease the production of ubiquinone-8 because of the downregulation of UbiE, one of the downstream enzymes involved in this biosynthetic pathway. The decrease in ubiquinone-8 biosynthesis leads to reduced aerobic respiration in *ΔhflKC* cells, probably due to low activity of cytochrome ubiquinol oxidases. This could be enhanced by their perturbed expression, including downregulation of the major *E. coli* cytochrome ubiquinol oxidase *bo_3_* (CyoABCD), which operates under high O_2_ conditions, and upregulation of the less efficient cytochrome ubiquinol oxidase *bd* (and CydAB), which normally operates under microaerobic conditions^31^.

This misregulation of cytochrome ubiquinol oxidases and several other respiration-related proteins could be largely explained by the activation of the two-component system ArcAB, which allows bacteria to adapt to changes in oxygen availability and activates the expression of genes involved in anaerobic respiration while inhibiting the expression of aerobic respiratory genes^36^. Its sensory kinase, ArcB, is normally repressed at high O_2_ by oxidized ubiquinone^35, 48^, but this repression appears to be alleviated in *ΔhflKC* cells due to the overall reduction in the ubiquinone levels, causing an aberrant activation of the ArcAB system. However, the levels of IspG and UbiE were affected by *ΔhflKC* deletion even in the absence of ArcB, confirming that the ArcAB system is downstream in this regulatory cascade.

In contrast to TB or minimal M9 medium, no growth defect was observed for *ΔhflKC* cells in LB containing yeast extract, even at high aeration. Compared to TB, changes in the levels of respiration-related proteins in LB were also limited to a smaller set of proteins, including IspG, UbiE, and both cytochrome oxidases. Possible explanations for this difference in growth could be the presence of isoprenoids or quinones in the yeast extract, which partially complement the effect of IspG and UbiE downregulation on ubiquinone biosynthesis and thus on respiratory activity, or a lower importance of respiration for *E. coli* growth in LB.

In addition to the respiratory proteins, the absence of the HflKC complex led to changes in the levels of a number of other proteins independent of IspG regulation. The most prominent group of these respiration-independent proteins are those involved in *E. coli* motility. The levels of all classes of motility proteins were reduced in the *ΔhflKC* strain, indicating that the underlying mechanism is based on changes in the levels or activity of an upstream regulator.

What is the cause of the severe reduction of IspG levels in *ΔhflKC* cells? Our results suggest that a decreased stability of IspG in the absence of the HflKC complex may be at least partly responsible for its lower level. This is most likely explained by increased degradation of IspG by FtsH in the absence of the HflKC complex, although this hypothesis remains to be proven. However, as the observed increase in IspG degradation was modest, other post-transcriptional regulatory mechanisms could not be ruled out.

Interestingly, although the FtsH-regulatory HflKC complex normally contains equal number of HflK and HflC subunits^22^, we observed a striking asymmetry in the effects of individual deletions of the *hflK* and *hflC* genes. While the loss of *hflK* causes phenotypes similar to the absence of the entire HflKC complex, deletion of the *hflC* gene alone has no apparent effect and only slightly enhances the phenotype of the *hflK* deletion. This observation is even more surprising considering that deletion of *hflC* causes a decrease in the level of HflK, as is frequently the case for the unassembled components of the heterooligomeric complexes^49, 50^. This implies that HflK alone, even at reduced protein levels, can largely carry out the function of the HflKC complex. Although the overall structures of HflK and HflC are similar, HflK has an additional C-terminal extension that resides inside the HflKC complex and interacts with FtsH, indicating that HflK may be more important for the assembly of the HflKC-FtsH complex and for FtsH regulation^22, 23^.

Although HflK and HflC are phylogenetically distant from eukaryotic prohibitins PHB1 and PHB2, the PHB1-PHB2 complex also forms a ring-like heterooligomer in the mitochondrial membrane that regulates the activity of an AFG3L2 AAA+ metalloprotease homologous to FtsH^22^ ^24, 25^. Notably, the PHB1-PHB2 complex is important for respiratory activity in human cells^15^ and associates with respiratory proteins^51, 52^. It has therefore been proposed to be involved in the assembly of respiratory complexes^14, 53, 54^, but the relationship between such putative chaperone activity and the control of the associated protease by the PHB1-PHB2 complex remained unclear. Our results demonstrate a different mechanism of regulation of respiratory activity by the bacterial analog of this complex, through control of ubiquinone biosynthesis. Although the relevance of this mechanism for eukaryotes remains to be investigated, the structural and functional similarity of the HflKC and PHB1-PHB2 complexes suggests that a similar mechanism may operate in mitochondria.

## ACKNOWLEDGMENTS

We thank Jörg Kahnt, Peter Claus, and Silvia Gonzalez Sierra for their technical assistance with proteomics, lipidomics, and flow cytometry measurements, respectively. We thank Seigo Shima for his help with anaerobic cultivation, Andreas Brune and Evgenii Protasov for help with oxygen consumption measurements, and Jorina Eckersberg for help with bacterial growth measurements. We thank Jing Yuan for her insightful discussions. This research was funded by the Max Planck Society and the IMPRS-µLife graduate program.

## AUTHOR CONTRIBUTIONS

M.I.P.L. and V.S. conceived the study; M.I.P.L., P.L., N.P., and T.G. performed the measurements; M.I.P.L., P.L., G.A., N.P., T.G., and H.L. analyzed the data; M.I.P.L. and V.S. wrote the paper with input from all other authors.

## STAR METHODS

### KEY RESOURCES TABLE

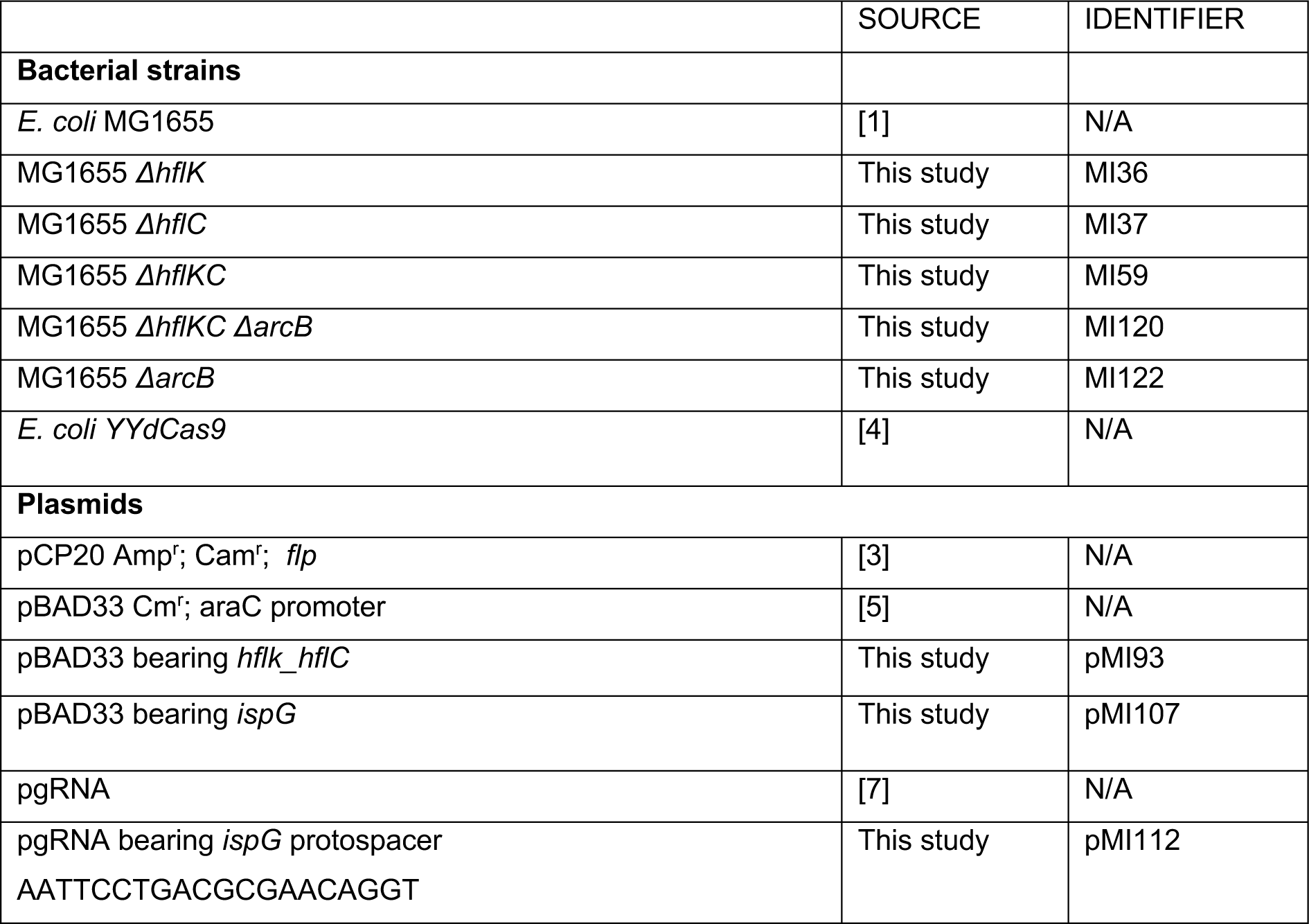

### RESOURCE AVAILABILITY

#### Lead contact

Further information and requests for resources and reagents should be directed to and will be fulfilled by the lead contact, Victor Sourjik.

#### Materials availability

All plasmids and strains generated in this study are available from the lead contact without restriction.

### METHOD DETAILS

#### Bacterial strains, plasmids, and growth conditions

*Escherichia coli* K-12 MG1655^1^ was used as the wild-type (WT) strain in this study. *ΔhflK* and *ΔhflC* gene deletions were constructed using P1 transduction from the Keio collection strains (JW 4132 and JW 4133, respectively). *ΔhflKC, ΔhflKC ΔarcB,* and *ΔarcB* strains were constructed using lambda red recombination as described previously^2^. Kanamycin cassettes were flipped out using FLP-FLP recombination target (FRT) recombination^3^. All knockout constructs were verified by PCR. *E. coli* YYdCas9 derived from *E. coli* K-12 (BW25993) was used as a background strain to construct *ispG* knockdown as described previously^4^. Plasmid expression vectors carrying *hflK-hflC* and *ispG* genes were constructed by amplifying DNA fragments from the MG1655 genome by PCR using Q5 high-fidelity DNA polymerase and cloned into pBAD33^5^ using Gibson assembly^6^. All strains and plasmids are listed in the Table below.

Strains were grown in LB medium (10 g tryptone, 10 g NaCl, and 5 g yeast extract per liter), TB medium (10 g tryptone and 5 g NaCl per liter), TB supplemented with 0.4% of glucose or M9 minimal medium with glucose as sole carbon source (5 g L-1). M9 medium was composed by (per liter): 7.52 g Na_2_HPO_4_ 2H_2_O, 5 g KH_2_PO_4_, 1.5 g (NH_4_)2SO_4_, 0.5 g NaCl. The following components were sterilized separately and then added (per liter of final medium): 1 mL 0.1 M CaCl_2_, 1 mL 1 M MgSO_4_, 0.6 mL 0.1 M FeCl_3_, 2 mL 1.4 mM thiamine HCl, and 10 mL trace salts solution. The trace salts solution contained (per liter): 180 mg ZnSO_4_ 7H_2_O, 120 mg CuCl_2_ 2H_2_O, 120 mg MnSO_4_ H_2_O and 180 mg CoCl_2_ 6H_2_O. Antibiotics (Kanamycin 50 µg/ml, Ampicillin 100 µg/ml, Chloramphenicol 34 µg/ml) and inducers of expression were added where necessary.

For all measurements, overnight cultures were diluted 1:100 in 50 ml fresh media and grown in 100 ml flasks at 37°C on an orbital shaker at indicated shaking rates (100 rpm, 220 rpm, or 300 rpm). For anaerobic growth, sealed flasks where oxygen was replaced with nitrogen were used.

#### Construction of *ispG* knockdown

Different protospacers designed along *ispG* gene were cloned in the plasmid vector pgRNA^7^*. Plasmids were then* transformed into *E. coli* YYdCas9. Expression of dCas9 was induced with 0.02 µM aTC (**anhydrotetracycline)**. The protospacer with the strongest effect of *ispG* knockdown on growth (AATTCCTGACGCGAACAGGT; pMI112) was selected for further experiments.

#### Total cell proteomics

Cultures were grown until OD_600_ of 0.4 for aerobic and 0.15 for anaerobic growth. Biomass was adjusted to OD_600_ = 3 in 1ml to have an equal amount of cells per sample. Pellets were washed twice with ice-cold 1X PBS (phosphate-buffered saline) and stored at -80°C.

For protein extraction, cell pellets were dissolved in 300 ul of 2% sodium-lauroyl sarcosinate (SLS) and 100 mM ammonium bicarbonate. Cells were lysed by incubation at 90°C for 15 minutes and subsequent sonication (Vial Tweeter, Hielscher) with 80% amplitude for 30 seconds. Cell lysates were reduced by adding 5 mM (final concentration) Tris(2-caboxyethyl)phosphine and incubating at 95°C for 15 minutes followed by alkylation (10 mM iodoacetamide final concentration, 30 minutes at 25°C).

The amount of extracted proteins was measured using BCA protein assay (Thermo Fisher Scientific). 50 µg total protein was then digested with 1 µg trypsin (Promega) overnight at 30 °C in the presence of 0.5% SLS. Following digestion, SLS was precipitated with trifluoroacetic acid (TFA, 1.5% final concentration) and peptides were purified using Chromabond C18 microspin columns (Macherey-Nagel). Acidified peptides were loaded on spin columns equilibrated with 400 µL acetonitrile and then 400 µL 0.15% TFA. After peptide loading, a washing step with 0.15% TFA was performed, followed by elution using 400 µL 50% acetonitrile. Eluted peptides were then dried by vacuum concentrator and reconstituted in 0.15% TFA.

Peptide mixtures were analyzed using liquid chromatography-mass spectrometry using an Ultimate RSLC nano connected to a Q-Exactive Plus mass spectrometer (both Thermo Scientific) as reported previously^8^. In short, peptides were separated using a gradient from 96 % solvent A (0.15% formic acid) and 4 % solvent B (99,85 % acetonitrile, 0.15 % formic acid) to 30 % solvent B over 90 or 120 minutes at a flow rate of 300 nL/min. MS data was acquired with the following settings: 1 MS scan at a resolution of 70,000 with 50 ms max. ion injection fill time, MS/MS at 17,500 scans of the 10 most intense ions with 50 ms maximum fill time. The data was further analyzed using either Progenesis (Waters) or MaxQuant in standard settings^9^ using an *E.coli* uniprot database. Follow up data analysis and data visualization was done with SafeQuant^10^ (available under https://github.com/eahrne/SafeQuant), Perseus^11^ and Rstudio software. Due to an instrumental upgrade a part of the total proteome samples were analyzed on an Exploris 480 connected to an Ultimate 3000 RSLC nano. The LC peptide separating gradient was reduced to 60 min (6-35% solvent B). The MS data was acquired in data independent acquisition mode (DIA) using 45 windows with an isolation window of 14 mz with 1 m/z overlap (see also^12^). MS scan resolution was set to 120,000 (MS1) and 15,000 (DIA) with a scan range of 350-1400 m/z (MS1) and 320-950 precursor mass range (DIA). AGC target settings were 300 % (MS1), and 3000 % (DIA) with a maximum ion injection time of 50 ms (MS1) and 22 ms (DIA). DIA data were analyzed using DIA-NN version 1.8^13^ and an *E.coli* protein database. Full tryptic digest was allowed with two missed cleavage sites, and oxidized methionines and carbamidomethylated cysteins. Match between runs and remove likely interferences were enabled. The neural network classifier was set to the single-pass mode, and protein inference was based on genes. Quantification strategy was set to any LC (high accuracy). Cross-run normalization was set to RT-dependent. Library generation was set to smart profiling. DIA-NN outputs were further evaluated using SafeQuant and data visualized in Perseus.

#### Stability measurements for IspG

Cultures were grown in LB and TB media and 220 rpm and 100 rpm, respectively. Samples were collected at OD_600_ = 0.6, and the biomass was adjusted to OD_600_ = 3 in 1 ml. Subsequently, chloramphenicol was added to the final concentration of 200 µg/ml, and the cultures were further incubated under the same conditions. Samples were collected after 30 and 60 minutes of incubation. All samples were washed twice with 1X PBS, and pellets were stored at -80°C until proceeding with the protein extraction and analysis by mass spectrometry as described above.

#### Quantification of ME-cPP and ATP measurements

Cultures were grown in M9 minimal medium supplemented with glucose at 220 rpm. Cells were grown to an OD_600_ = 0.4 - 0.5, this preculture was used to inoculate cultures at a final volume of 10 ml M9 glucose minimal medium and starting OD_600_ = 0.05, which were allowed to grow until OD_600_ = 0.5. Biomass of OD_600_ = 0.8 was applied on filter disc (PVDF Membranes: 0.45μ pore size) and immediately transferred into 1ml acetonitrile: methanol: water (40:40:20 (v/v)) kept at -20°C. Samples were incubated for 30 minutes at -20°C. After that time, 500ul of the samples were transferred into a 1.5ml tube at -20°C and centrifuged at -9°C and >13.000 rpm for 15 minutes. 350 μl of supernatant was transferred to new Eppendorf tubes and stored at -80°C until LCMS analysis. 15 µl of each sample was mixed with 15 µl of ^13^C-labeled internal standard. Analysis of target metabolites was performed with an Agilent 6495 triple quadrupole mass spectrometer (Agilent Technologies) and an Agilent 1290 Infinity II UHPLC system (Agilent Technologies) as described previously^14^. The temperature of the column oven was 30 °C, and the injection volume was 3 μl. LC solvents A were water with 10 mM ammonium formate and 0.1% formic acid (v/v) (for acidic conditions); and water with 10 mM ammonium carbonate and 0.2% ammonium hydroxide (for basic conditions). LC solvents B were acetonitrile with 0.1% formic acid (v/v) for acidic conditions and acetonitrile without additive for basic conditions. LC columns were an Acquity BEH Amide (30 × 2.1 mm, 1.7 µm) for acidic conditions, and an iHILIC-Fusion(P) (50 × 2.1 mm, 5 µm) for basic conditions. The gradient for basic and acidic conditions was: 0 min 90% B; 1.3 minutes 40% B; 1.5 minutes 40% B; 1.7 minutes 90% B; 2 minutes 90% B. Quantification of metabolite concentrations was based on the ratio of ^12^C and ^13^C peak heights.

#### Quantification of ubiquinone-8 and ubiquinol-8

Cultures were grown in TB at 200 rpm until OD_600_ = 0.4 – 0.8. Biomass was adjusted to OD_600_ = 5 in 1ml. Cells were collected by centrifugation and washed twice with 1X PBS. Pellet samples were dissolved in a mixture of 150 µl of chloroform, 300 µl of methanol, and 120 µl of water; followed by shaking for 10 minutes at 4°C. Afterward, 150 µl of chloroform and 150 µl of 0.85 % KCL were added. Samples were centrifuged for 10 minutes at max g at 4°C. The lipid phase was transferred to new tubes and dried out with nitrogen. The relative quantification and annotation of lipids were performed by using HRES-LC-MS/MS. The chromatographic separation was performed using a Acquity Premier CSH C18 column (2.1 × 100 mm, 1.7 μm particle size, Waters, Milford, USA) a constant flow rate of 0.3 ml/min with mobile phase A being 10mm Ammonium Formate in 6:4 Acetonitrile:water and phase B being 9:1 Isopropanol:Acetonitrile (Honeywell, Morristown, New Jersey, USA) at 40° C. The injection volume was 5 µl. The mobile phase profile consisted of the following steps and linear gradients: 0 – 1.5 min constant at 37 % B; 1.5 – 4 min from 37 to 45% B; 4 – 5 min from 45 to 52% B; 5 – 8 min from 52 to 58 % B; 8 - 11 min from 58 to 66 % B; 11 - 14 min from 66 to 70 % B; 11 - 14 min from 66 - 70 % B; 14 - 18 min from 70 to 75 % B; 18 - 20 min from 75 to 98 % B; 20 - 25 min constant at 98 % B; 25 – 25.1 min from 98 to 37 % B; 25.1 – 30 min constant at 37 % B.

For the measurement, a Thermo Scientific ID-X Orbitrap mass spectrometer was used. Ionisation was performed using a high temperature electro spray ion source at a static spray voltage of 3500 V (positive) and a static spray voltage of 2800 V (negative), Sheath gas at 50 (Arb), Auxilary Gas at 10 (Arb), and Ion transfer tube and Vaporizer at 325 and 300°C. Data dependent MS2 Measurement were conducted applying an orbitrap mass resolution of 120 000 using quadrupole isolation in a mass range of 200 – 2000 and combining it with a high energy collision dissociation (HCD). HCD was performed on the ten most abundant ions per scan with a relative collision energy of 25 %. Fragments were detected using the orbitrap mass analyser at a predefined mass resolution of 15 000. Dynamic exclusion with and exclusion duration of 5 seconds after 1 scan with a mass tolerance of 10 ppm was used to increase coverage.

Compound Discoverer 3.3 (Thermo-Fisher Scientific) was used for lipid annotation by matching accurate mass and MS2 spectra against the MS/MS library MS-DIAL LipidBlast (version 68). In addition, two customized in-house libraries were used for the annotation of the target analytes Ubiquinone-8 and Ubiquinol-8, and a set of eight lipids that served as internal standards. For the semi-quantitative comparison of lipid abundance, annotated peaks were integrated using Compound Discoverer 3.3 (Thermo Scientific) and normalization by the default method provided by Compound Discoverer 3.3 and further processed by the statistical tools described elsewhere.

Ubiquinol annotation was done employing Compound Discoverer 3.3 (CD) using a customized CD workflow and matching the metabolic features against three different data libraries. The majority of lipids were matched against the MS-Dial LipidBlast library (version68). In addition two customized in-house libraries were used. The “IS-List.massList” contained the names of the 8 lipids that were used as internal standards (LPE 13:0, PE 40:8, PG 40:8, CL 56:4, Cer 22:1;2, HexCer 26:1;2 and SM 24:1;2) and the “targetedCompounds.massList” contained the ammonium adduct of the ubiquinol-8 and ubiquinone-8 (CoQ8). The library focus for the targeted analytes was created by the in-house MS/MS measured spectra from previous runs and the library focus in the internal standards was created base the theoretical mass calculated by the elemental formula.

#### Measurements of oxygen consumption

Strains were grown in TB at 37°C and 220 rpm until OD_600_ = 0.4. Biomass was adjusted to an OD_600_ = 1 in 5ml. Cultures were centrifuged and fresh TB medium was added. Samples were transferred to a glass tube that contained an oxygen sensor spot PSt3-YAU-D5-YOP (PresSens, precision sensing). Sample tubes were under vortex for 1 minute to achieve maximum oxygenation; then shaking was stopped and oxygen consumption was measured via the oxygen spot with a fiber optic transmitter.

#### Measurement of reactive oxygen species (ROS)

The dichlorodihydrofluorescein (DCF) fluorescent probe by **Abcam (ab113851 Kit) was used to measure r**eactive oxygen species. Strains were grown in TB 37°C and 220 rpm until OD_600_ = 0.4. Biomass was adjusted to have OD_600_ = 0.4 in 1 ml. Samples were transferred to a 1.5 ml Eppendorf tube where DCF probe was added to have a final concentration of 20 µM. Samples were gently mixed by inversion, followed by dark incubation for 30 minutes in the dark at 37°C. Fluorescence was analyzed by flow cytometry at 485nm. Treatment with 0.5 mM of hydrogen peroxide (H_2_O_2_) was used as a positive control for elevated ROS levels.

#### Characterization of membrane potential (MP)

BacLight Bacterial Membrane Potential kit (B34950 Molecular Probes) was used to measure the membrane potential. Strains were grown in TB at 37°C and 220 rpm. All samples were diluted in 1X PBS and biomass was adjusted to have OD_600_ = 0.4 in 1ml. Samples were transferred to a 1.5 ml Eppendorf tube where DiOC_2_(3) was added to a final concentration of 0.03 mM. Samples were gently mixed by inversion, followed by incubation for 15 minutes at 37°C in the dark. WT treated with 40 µM of dinitrophenol (DNP) was used as a negative control. Flow cytometry was used to measure the fluorescence of red (670 nm) and green (510 nm) channels of DiOC_2_(3). Excitation at 488 nm was used and fluorescence was measured through a 530-nm bandpass filter. MP was characterized by the ratio of the red and green fluorescence according to the manufacturer’s instructions.

#### RNA extraction and real-time PCR (RT-PCR)

Strains were grown in LB medium at 220 rpm until OD_600_ = 0.4. Cultures were concentrated to have OD_600_ = 1 in 1ml. After centrifugation pellets were washed twice with cold water and stored at -80°C. Frozen pellets were resuspended in 800 μl of lysis buffer (2 % SDS and 4 mM EDTA) and boiled for 2 minutes at 90°C. Subsequently, 800 μl of TRIzol was added and incubated at room temperature for 5 minutes. To the mixture, 200 μl of phenol:chloroform was added, vortexed for 30 seconds, and incubated for 10 minutes. Samples were then centrifuged at 13,000 x g and 4°C for 10 minutes to separate the phases. The upper aqueous phase containing RNA was transferred to a new tube containing 500 μl of isopropanol for RNA precipitation, which was carried out overnight at -20°C. The following day, samples were centrifuged at 13,000 x g and 4°C for 30 minutes, and the supernatants were discarded. RNA pellets were washed twice with 70% ethanol, air-dried, and resuspended in 50 μl of nuclease-free water to proceed with DNase treatment. After that, samples were stored at -80°C.

The RT-PCR reactions were performed as described in KAPA SYBR FATS one-step qRT-PCR master mix 2X Kit (KR0393) using 2 µl of 10 ng/μl RNA sample. Primers used for IspG were GTATTTACGTTGGGAATGTGCCG and GATATCAGCGCCAACGCGTTC. Housekeeping gene ssrA was used as a control with primers ATTCTGGATTCGACGGGATT and AGTTTTCGTCGTTTGCGACT.

### QUANTIFICATION AND STATISTICAL ANALYSIS

Details on the number of replicates, the sample sizes as well as the value and meaning of n are included in the figure legends.

## SUPPLEMENTAL INFORMATION

Supplemental figures: Figures S1–S9

Supplemental tables: Tables S1-S9

