## Supplemental Figures for "The SPFH complex HflK-HflC regulates aerobic respiration in bacteria"

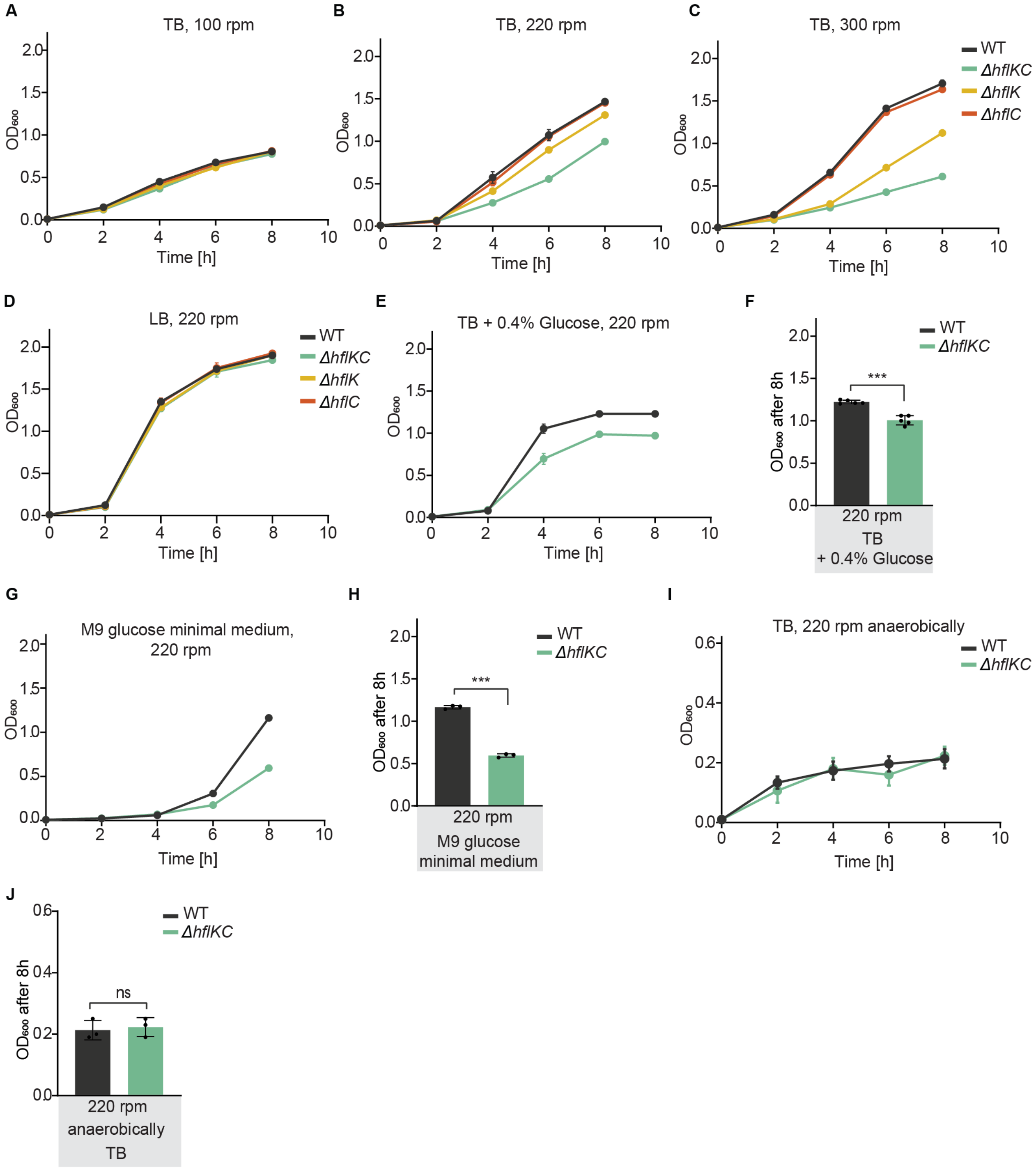


**Figure S1. Growth of *ΔhflKC, ΔhflK,* and *ΔhflC* strains under different conditions. Related to Figure 1.**

(A - C) Growth of *E. coli ΔhflKC, ΔhflK,* and *ΔhflC* strains and corresponding wild-type (WT) in TB medium in an orbital shaker at 100 rpm (A), 220 rpm (B), or 300 rpm (C), quantified by optical density at 600 nm ($OD_{600}$). Data represent the mean of three independent cultures, different from the experiment shown in Figure 1A-C. Error bars indicate the standard deviation (SD).

(D) Growth of the indicated strains in LB medium in an orbital shaker at 220 rpm. Data represent the mean and SD of three independent cultures, different from the experiment shown in Figure 1G.

(E, F) Growth of *ΔhflKC* and WT strains in TB supplemented with 0.4% glucose at 220 rpm (E) and corresponding final $OD_{600}$ after 8 h of growth (F). Data in (E) represent the mean and SD of three independent cultures. Data in (F) represent the mean and SD of independent c indicated by dots, grown in three different experiments.
(G, H) Growth of *ΔhflKC* and WT strains in M9 glucose minimal medium (G) and corresponding final $OD_{600}$ after 8 h of growth (H). Data represent the mean and SD of three independent cultures grown in the same representative experiment.
(I, J) Growth of *ΔhflKC* and WT strains in TB at 220 rpm under anaerobic conditions (I) and corresponding final $OD_{600}$ after 8 h of growth (J). Data represent the mean and SD of three independent cultures grown in the same representative experiment.
Indicated differences between samples: ***$p<0.001$ or not significant (ns) by unpaired *t*-test.

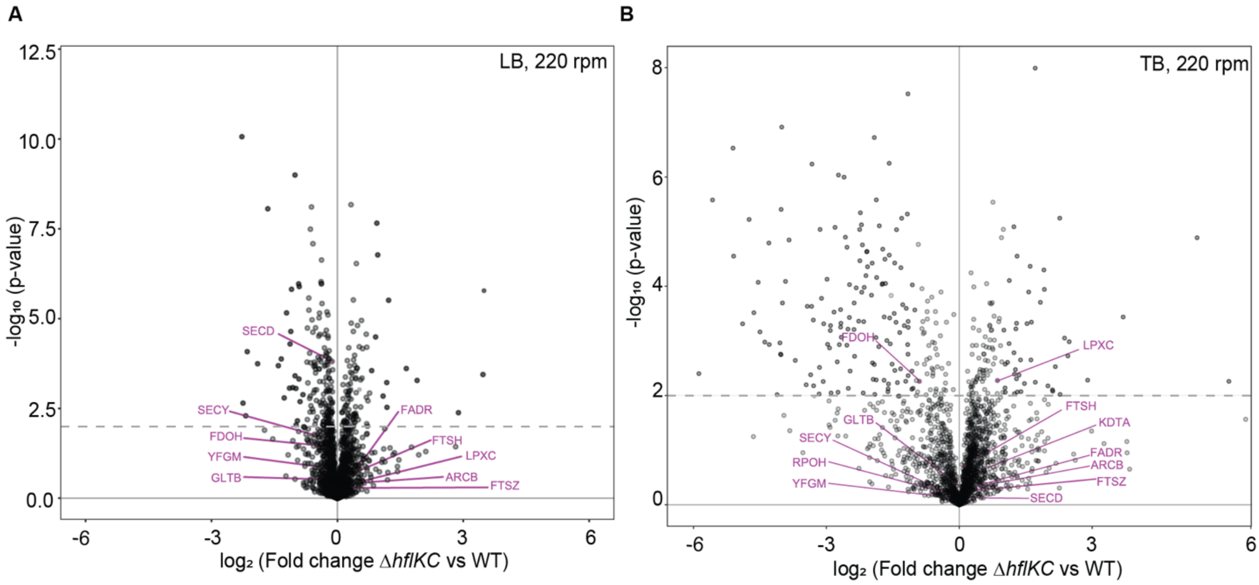


**Figure S2. Comparison of abundances of known FtsH substrates between *ΔhflKC* and wild-type cells. Related to Figure 2.**

(A, B) Differences in protein levels between *ΔhflKC* and wild-type (WT) strains grow in LB (A) or TB (B) at 220 rpm. Data are the same as in Figures 2A and 2B, but with known FtsH substrates and FtsH itself labeled in purple. Data are for six (LB) or three (TB) independent cultures.

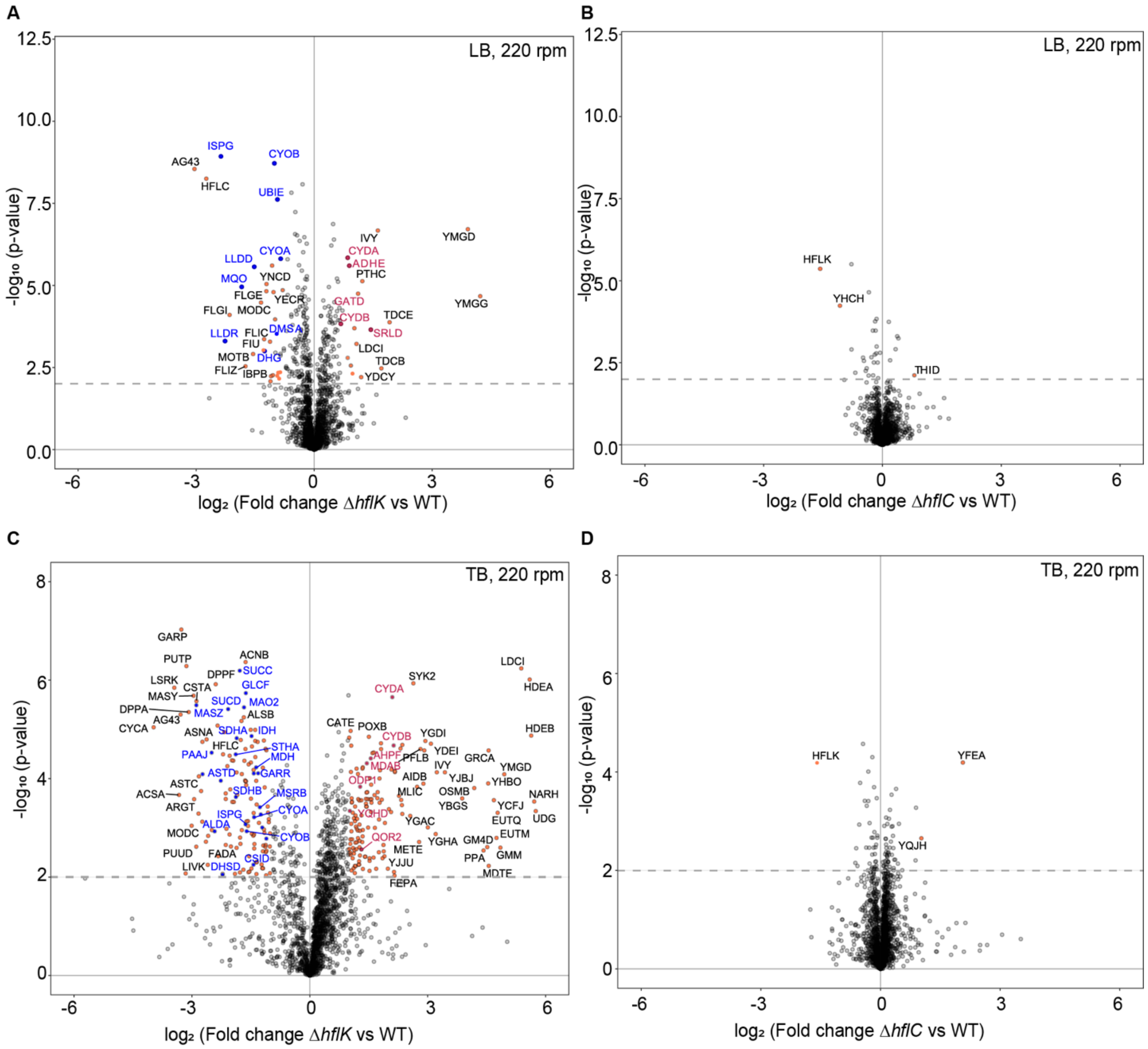


**Figure S3. Changes in protein abundance in the absence of HflK or HflC proteins. Related to Figure 2.**

(A-D) Differences in protein levels between the *ΔhflK* (A,C) and *ΔhflC* (B,D) strains and the wild-type (WT), for cultures grown for 4 h at 220 rpm in LB (A,B) or TB (C,D). Data for LB are for six independent cultures; data for TB are for three independent cultures. Proteins with differences in expression that were considered significant (see also Table 1, Table S1 and Table S2) are labeled, with respiration-related proteins highlighted in either blue (downregulated) or red (upregulated).

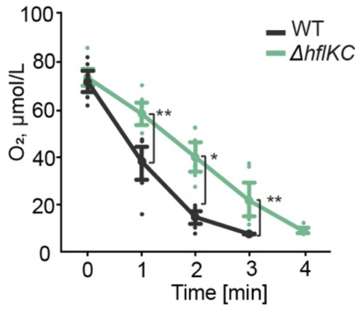


**Figure S4. Oxygen consumption by *ΔhflKC* and wild-type strains. Related to Figure 3.**

A different biological replica of the experiment shown in Figure 3E. *ΔhflKC* and wild-type (WT) were grown in TB at 220 rpm, resuspended in fresh TB, and changes in the levels of dissolved oxygen were quantified. The lines represent the average of eight independent measurements for one culture, with error bars indicating standard deviation. *$p<0.05$, **$p<0.01$ by unpaired *t*-test.

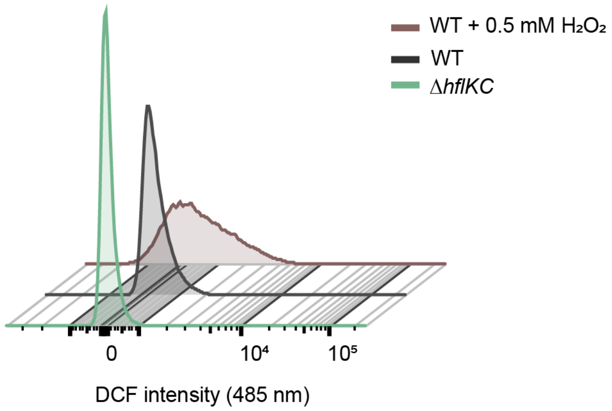


**Figure S5. Levels of ROS in wild-type and *ΔhflKC* cells grown in TB at 220 rpm. Related to Figure 3.**

Cells were grown until the early exponential phase and biomass was adjusted to $OD_{600}$ equal to 0.4 in 1 ml. Dichlorodihydrofluorescein (DCF) probe was added to the samples following incubation for 30 min in the dark. Fluorescence (485nm) was analyzed via flow cytometry. Treatment with hydrogen peroxide ($H_2O_2$) was used as a positive control for elevated ROS levels. The figure is an illustration of one representative measurement.

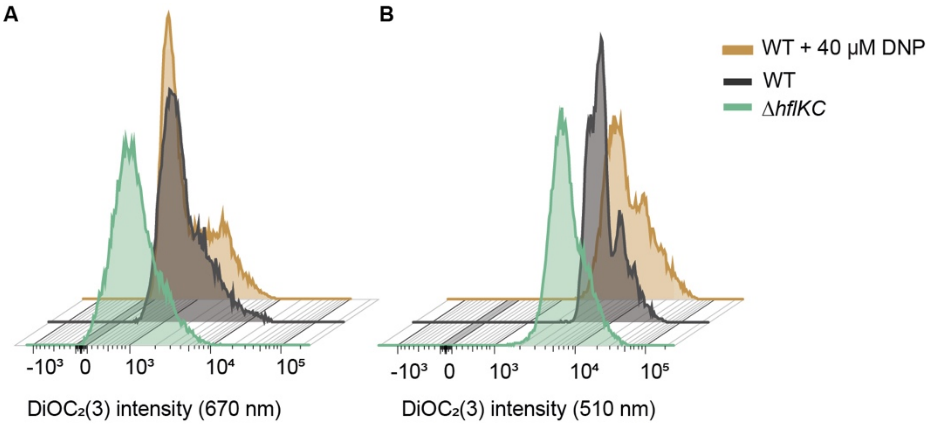


**Figure S6: Membrane potential in wild-type and *ΔhflKC* cells grown in TB at 220 rpm. Related to Figure 3.**

Cells were grown until the early exponential phase and biomass was adjusted to $OD_{600}$ equal to 0.4 in 1 ml. $DiOC_2(3)$ probe was added to the samples following incubation for 15 min in the dark. Fluorescence in the red (670 nm) and green (510 nm) was analyzed via flow cytometry. The protonophore dinitrophenol (DNP) that dissipates the proton gradient across the cytoplasmic membrane was used as a control. The figure is an illustration of one representative measurement.

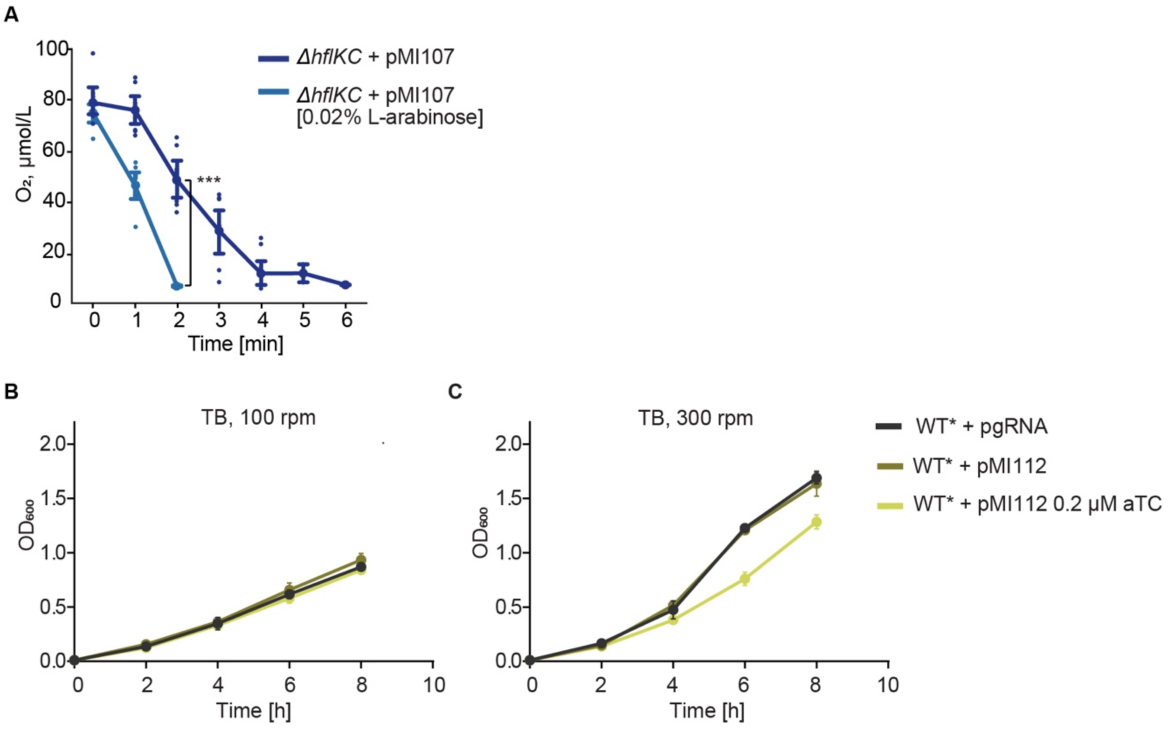


**Figure S7. Impact of IspG expression on oxygen consumption and growth. Related to Figure 4.**
(A) A different biological replicate of the experiment shown in Figure 4C. Oxygen consumption by the *ΔhflKC* strain carrying the IspG expression construct pMI107. Cells were grown in TB medium at 220 rpm, with either no or 0.02% L-arabinose induction, as indicated. The lines represent the average of eight measurements for one biological replicate. Error bars indicate the SD. ***$p<0.001$ by unpaired *t*-test.
(B, C) Growth of *E. coli* YYdCas9 (WT*) strain carrying either the control pgRNA ter or pgRNA *ispG* (pMI112). When indicated, dCas9 was induced by adding 0.2 µM of anhydrotetracycline (aTC). Cells were grown in TB medium at 100 rpm (B) and 300 rpm (C). Data represent the mean value and standard deviation (SD) for three independent cultures.

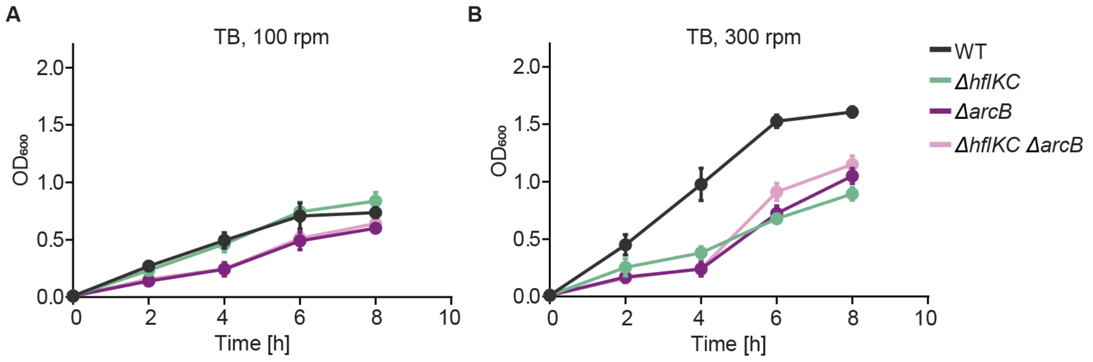


**Figure S8. Dependence of *ΔhflKC* deletion growth effect on *ΔarcB* background. Related to Figure 5.**

(A, B) Growth of the wild-type (WT), *ΔhflKC, ΔarcB,* and *ΔhflKC ΔarcB* strains in TB medium in a rotary shaker at 100 rpm (A), 300 rpm (B). Data represent the mean of three independent cultures (±SD).

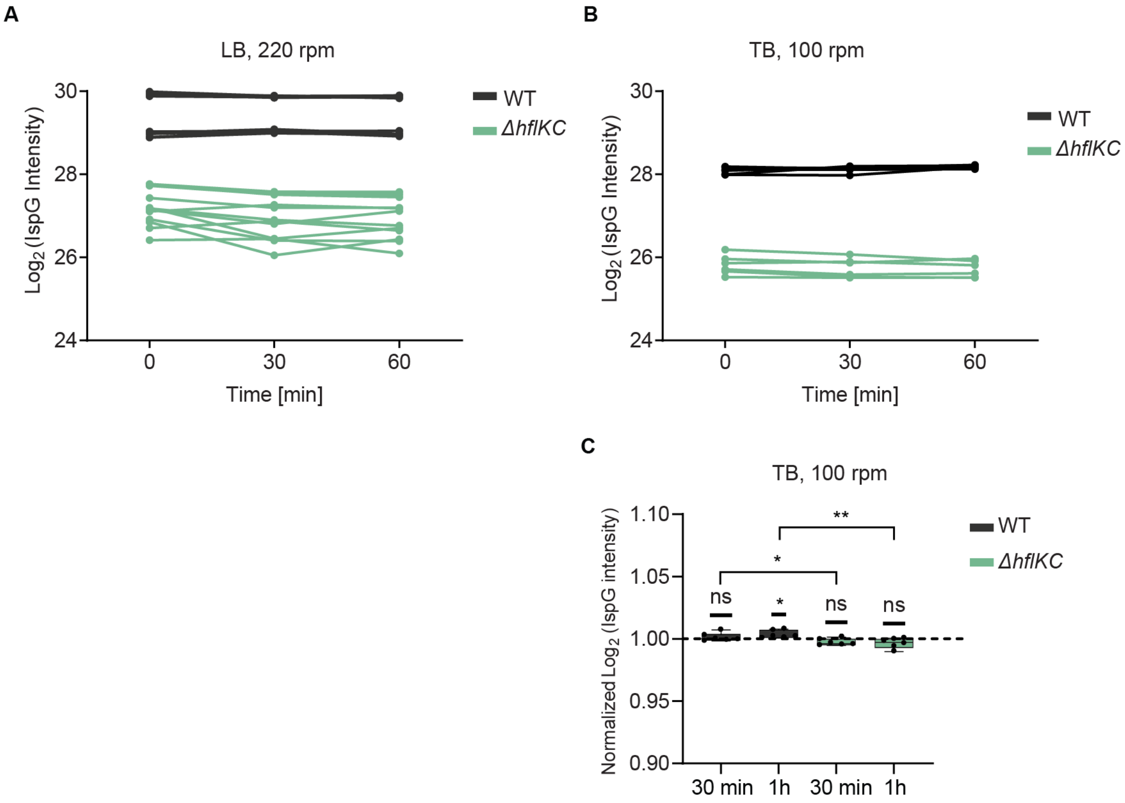


**Figure S9: Stability of IspG in the wild-type and *ΔhflKC* strains. Related to Figure 6.**

(A) The abundance of IspG was determined using proteomics. Strains were incubated with chloramphenicol in LB at 220 rpm for 30 minutes or 1 hour. The abundance of IspG is represented by $Log_2$ intensity. Data represent the mean and SD of two independent experiments with six independent cultures each.

(B, C) The abundance of IspG was determined as in panel A, however, the strains were grown in TB at 100 rpm (B). Abundance of IspG was normalized to the initial time point (C). Data represent the mean and SD of two independent experiments with six independent cultures each. Significance of difference from the value of 1 by one sample *t*-test is indicated, as well as significance of difference between wild-type (WT) and *ΔhflKC* values at the same time point (indicated by brackets): *$p < 0.05$, **$p < 0.01$, and ns = not significant.
