## Supplemental Tables for "The SPFH complex HflK-HflC regulates aerobic respiration in bacteria"

**Table S1. Other proteins showing significant differences between *ΔhflKC, ΔhflK* or *ΔhflC* and wild-type strains during growth in LB at 220 rpm**

| | | $Log_2$ (Fold change) | | |
|---|---|---|---|---|
| Protein | Function | *ΔhflKC* vs WT | *ΔhflK* vs WT | *ΔhflC* vs WT |
| YEJG | Unknown function | -2.25 | -1.27 | -0.15 |
| YIGI | Putative thioesterase | -2.18 | -1.23 | -0.14 |
| IBPB | Small heat shock protein | -1.74 | -1.10 | -0.75 |
| MODC | Molybdate ABC transporter ATP binding subunit | -1.41 | -1.35 | -0.07 |
| MOTB | Flagellar rotation | -1.27 | -1.55 | -0.19 |
| YNCD | Pyrroloquinoline quinone TonB-dependent | -1.21 | -1.21 | -0.11 |
| TSR | Methyl-accepting chemotaxis protein I | -1.14 | -1.12 | -0.20 |
| EFEO | Ferrous iron transport system | -1.11 | -0.99 | -0.12 |
| KGTP | Alpha-ketoglutarate:H(+) symporter | -1.09 | -1.07 | -0.35 |
| FIU | Iron transport | -1.06 | -1.29 | -0.06 |
| FLGI | Flagellar P-ring protein | -1.06 | -2.15 | -0.12 |
| FLHE | Flagellar protein | -1.01 | -0.86 | -0.19 |
| FLIC | Flagellar filament structural protein | -0.99 | -1.27 | -0.21 |
| CHEY | Chemotaxis protein | -0.98 | -0.84 | -0.39 |
| TAP | Methyl-accepting chemotaxis protein IV | -0.95 | -1.11 | -0.42 |
| FLIZ | DNA-binding transcriptional regulator | -0.92 | -1.75 | 0.19 |
| OMPF | Outer membrane porin F | -0.92 | -0.86 | -0.22 |
| SMG | Uncharacterized protein | -0.89 | -0.12 | -0.24 |
| YCGR | Flagellar brake protein | -0.82 | -0.86 | -0.07 |
| YHCH | N-acetylneuraminate anomerase | -0.80 | -0.35 | -1.07 |
| MOTA | Motility protein A | -0.80 | -1.05 | 0.10 |
| FLGN | Flagellar biosynthesis protein | -0.79 | -0.99 | 0.09 |
| CHEA | Chemotaxis protein | -0.77 | -0.91 | 0.03 |
| CHEW | Chemotaxis protein | -0.75 | -0.89 | -0.06 |
| FLGE | Flagellar hook protein | -0.70 | -1.22 | -0.16 |
| DEAD | ATP-dependent RNA helicase | -0.58 | -0.80 | -0.01 |
| YECR | Unknown function | -0.58 | -1.03 | -0.05 |
| YMGD | Unknown function | 3.49 | 3.91 | 0.39 |
| YMGG | Unknown function | 3.47 | 4.22 | 0.26 |
| YKGG | Unknown function | 2.88 | 1.10 | 0.50 |
| TDCB | Catabolic threonine dehydratase | 1.90 | 1.71 | 0.63 |
| TDCE | Formate C-acetyltransferase | 1.64 | 1.92 | 0.44 |
| IVY | Inhibitor of vertebrate lysozyme | 1.22 | 1.62 | 0.00 |
| LDCI | Lysine decarboxylase 1 | 1.18 | 1.08 | 0.41 |
| YDCY | Unknown function | 1.13 | 1.19 | 0.94 |

| | | | | |
|---|---|---|---|---|
| YBHB | Putative kinase inhibitor | 1.09 | 0.93 | 0.43 |
| PTHA | Glucitol/sorbitol transport | 0.77 | 1.02 | -0.28 |
| PTHB | Glucitol/sorbitol transport | 0.72 | 0.97 | -0.44 |
| PTHC | Glucitol/sorbitol transport | 0.71 | 1.22 | -0.43 |
| TNAA | Tryptophanase | 0.62 | 0.85 | -0.15 |
| AG43 | Biofilm formation, Antigen 43 | 0.37 | -3.04 | -0.05 |
| THID | Hydroxymethylpyrimidine kinase | 0.22 | 0.53 | 0.81 |
| FLGC | Flagellar basal-body rod protein | ND | -0.94 | -0.02 |

ND: not detected

Data are from Figures 2A and S3A, S3B. Differences with p-value <0.05 and $Log_2$ of fold change >0.8 were considered significant.

**Table S2. Proteins showing significant differences between *ΔhflKC, ΔhflK* or *ΔhflC* and wild-type strains during growth in TB at 220 rpm**

| | | $Log_2$ (Fold change) | | |
|---|---|---|---|---|
| Protein | Function | *ΔhflKC* vs WT | *ΔhflK* vs WT | *ΔhflC* vs WT |
| Respiratory proteins | | | | |
| PUTA | Oxidoreductase, proline dehydrogenase | -4.30 | -2.73 | -0.10 |
| CSID | Oxidoreductase, glutarate hydroxylase | -4.29 | -1.43 | -0.05 |
| PUUC | NADP/NAD-dependent aldehyde dehydrogenase | -4.02 | -2.97 | -0.39 |
| ASTD | Aldehyde dehydrogenase | -3.43 | -2.27 | -0.06 |
| MASZ | Transferases, glyoxylate transacetylase | -3.33 | -2.89 | 0.13 |
| GLCE | Glycolate dehydrogenase, putative FAD-binding subunit | -3.16 | -2.46 | 0.30 |
| GLCF | Glycolate dehydrogenase | -3.14 | -1.63 | 0.31 |
| GABD | Succinate-semialdehyde dehydrogenase | -2.76 | -1.11 | -0.02 |
| ISPG | Oxidoreductase involved in isoprenoid biosynthesis | -2.71 | -1.64 | -0.01 |
| CYSI | Sulfite reductase, hemoprotein subunit | -2.67 | 0.07 | -0.10 |
| ALDA | Lactaldehyde dehydrogenase A | -2.60 | -2.42 | -0.11 |
| DHSD | Succinate:quinone oxidoreductase | -2.48 | -2.22 | -0.02 |
| GARR | Oxidoreductase, tartronate semialdehyde reductase | -2.37 | -1.43 | 0.21 |
| FADH | Oxidoreductase, 2,4-dienoyl-CoA reductase | -2.36 | -1.25 | 0.00 |
| LHGO | L-2-hydroxyglutarate dehydrogenase | -2.33 | -1.82 | -0.14 |
| STHA | Oxidoreductase, NAD(P)+ transhydrogenase | -2.21 | -1.89 | -0.07 |
| SUCC | Succinyl-CoA synthetase subunit β | -2.09 | -1.78 | -0.08 |
| SUCD | Succinyl-CoA synthetase subunit α | -2.08 | -2.08 | -0.11 |
| SDHB | Succinate:quinone oxidoreductase | -1.98 | -1.88 | -0.08 |
| SDHA | Succinate:quinone oxidoreductase | -1.97 | -1.86 | -0.06 |
| PUUB | Gamma-glutamylputrescine oxidase | -1.96 | ND | ND |

| | | | | |
|---|---|---|---|---|
| CYSJ | Sulfite reductase, flavoprotein subunit | -1.95 | 1.20 | 0.14 |
| LLDD | L-lactate dehydrogenase | -1.92 | -1.30 | 0.11 |
| MAO2 | Malate dehydrogenase | -1.92 | -1.67 | -0.05 |
| GLCD | Glycolate dehydrogenase, putative FAD-linked subunit | -1.68 | -0.51 | 0.10 |
| ODO1 | Oxidoreductase, 2-oxoglutarate dehydrogenase | -1.67 | -1.31 | -0.04 |
| DADA | D-amino acid dehydrogenase | -1.64 | -1.21 | -0.06 |
| UBIB | Ubiquinone biosynthesis protein | -1.61 | 0.21 | 0.38 |
| MSRB | Oxidoreductases, methionine sulfoxide reductase | -1.54 | -1.27 | -0.10 |
| IDH | Oxidoreductase, isocitrate dehydrogenase | -1.47 | -1.48 | -0.07 |
| CYOA | Cytochrome bo(3) ubiquinol oxidase subunit 2 | -1.42 | -1.42 | -0.02 |
| MDH | Malate dehydrogenase | -1.35 | -1.37 | -0.11 |
| CYOB | Cytochrome bo(3) ubiquinol oxidase subunit 1 | -1.20 | -1.61 | 0.11 |
| MDAB | NADPH:quinone oxidoreductase | 2.17 | 1.45 | -0.08 |
| DHNA | Type II NADH:quinone oxidoreductase | 1.92 | 2.16 | -0.02 |
| CYDB | Cytochrome bd-I ubiquinol oxidase subunit 2 | 1.62 | 2.13 | 0.08 |
| CYDA | Cytochrome bd-I ubiquinol oxidase subunit 1 | 1.53 | 2.10 | -0.07 |
| QOR2 | Quinone reductase 2 | 1.31 | 1.31 | -0.21 |
| ODP1 | Oxidoreductase, pyruvate dehydrogenase | 1.30 | 1.28 | 0.09 |
| YQHD | NADPH-dependent aldehyde reductase | 1.29 | 1.02 | -0.18 |
| AHPF | Oxidoreductase, alkyl hydroperoxide reductase | 1.12 | 1.55 | 0.13 |
| Other proteins | | | | |
| PAAJ | β-ketoadipyl-CoA thiolase | -5.57 | -2.50 | -0.45 |
| XYLF | Xylose ABC transporter periplasmic binding protein | -5.88 | -0.34 | -0.22 |
| ASTC | Succinylornithine aminotransferase | -5.11 | -2.82 | -0.11 |
| LSRR | DNA-binding transcriptional repressor | -4.89 | -0.96 | 0.02 |
| UGPB | Transport of sn-glycerol 3-phosphate | -4.64 | -2.12 | -0.16 |
| ACTP | Acetate transpor | -4.54 | -6.15 | -0.51 |
| POTF | Putrescine ABC transporter periplasmic binding protein | -4.50 | -0.82 | 0.11 |
| GARD | Galactarate dehydratase | -4.39 | -1.47 | 0.19 |
| PAAK | Phenylacetate-CoA ligase | -4.11 | -1.70 | -0.63 |
| DPPC | Dipeptide ABC transporter membrane subunit | -4.05 | -5.72 | -0.08 |
| GLCC | DNA-binding transcriptional dual regulator | -4.04 | -1.93 | -0.20 |
| FADI | Acetyl-CoA acyltransferase | -4.03 | -2.22 | -0.22 |
| ACSA | Acetyl-coenzyme A synthetase | -4.01 | -3.33 | -0.23 |
| LSRB | Autoinducer-2 ABC transporter periplasmic binding | -3.99 | -2.14 | -0.44 |
| YDCI | DNA-binding transcriptional dual regulator | -3.92 | -1.89 | -0.19 |
| GLCG | Unknown function | -3.84 | -2.83 | 0.19 |
| YDCS | Putative ABC transporter periplasmic binding | -3.7 | -1.77 | -0.16 |
| FADJ | Unsaturated acyl-CoA hydratase | -3.46 | -1.97 | 0.12 |
| DPPF | Dipeptide ABC transporter ATP binding subunit | -3.36 | -2.39 | -0.07 |

| | | | | |
|---|---|---|---|---|
| LSRF | 3-hydroxy-2,4-pentadione 5-phosphate thiolase | -3.33 | -2.49 | -0.30 |
| FUMC | Fumarate hydratase class II | -3.30 | -2.12 | -0.07 |
| TNAB | Tryptophan:H+ symporter, transporter of tryptophan | -3.29 | -2.14 | -0.09 |
| YHHZ | Putative endonuclease YhhZ | -3.16 | ND | ND |
| PAAX | DNA-binding transcriptional repressor | -3.11 | -1.30 | -0.43 |
| PAAY | Phenylacetic acid degradation protein | -3.07 | -0.27 | 0.35 |
| GUDD | D-glucarate dehydratase | -3.06 | -1.43 | 0.20 |
| CSTA | Pyruvate transporter | -2.99 | -2.88 | -0.22 |
| ASPG2 | L-asparagine 2 | -2.97 | 1.87 | 0.41 |
| TRPGD | Anthranilate synthase subunit | -2.93 | ND | ND |
| GARL | 5-keto-4-deoxy-D-glucarate aldolase | -2.91 | -2.14 | 0.12 |
| TRPC | Indole-3-glycerol phosphate synthase | -2.91 | -1.85 | -0.31 |
| GLXK1 | Glycerate 2-kinase | -2.9 | -1.17 | -0.23 |
| PUTP | Sodium/proline transporter | -2.83 | -3.14 | -0.24 |
| ARGT | Lysine, arginine and ornithine transport | -2.81 | -2.94 | -0.11 |
| PUUR | DNA-binding transcriptional repressor | -2.74 | -2.11 | -0.15 |
| TRPB | Tryptophan synthase subunit beta | -2.73 | -0.47 | 0.03 |
| PAAE | Phenylacetyl-CoA 1,2-epoxidase, reductase subunit | -2.68 | ND | ND |
| TRG | Methyl-accepting chemotaxis protein | -2.68 | -2.04 | -0.18 |
| ASTB | N-succinylarginine dihydrolase | -2.67 | -1.76 | -0.39 |
| DPPA | Dipeptide ABC transporter periplasmic binding protein | -2.57 | -3.08 | -0.10 |
| PUUD | Amine and polyamine degradation | -2.54 | -2.89 | -0.43 |
| PAAC | Phenylacetyl-CoA 1,2-epoxidase, structural subunit | -2.54 | ND | ND |
| PUUE | 4-aminobutyrate aminotransferase | -2.51 | -1.1 | -0.28 |
| ARAF | Arabinose ABC transporter periplasmic | -2.51 | -1.19 | -0.27 |
| GABT | 4-aminobutyrate aminotransferase | -2.48 | -1.90 | -0.08 |
| CYCA | D-serine/D-alanine/glycine transporter | -2.46 | -3.97 | -0.15 |
| ASTA | Arginine N-succinyltransferase | -2.38 | -0.34 | -0.13 |
| OMPF | Outer membrane porin F | -2.27 | -2.1 | -0.10 |
| GLNH | L-glutamine ABC transporter periplasmic binding protein | -2.25 | -1.95 | -0.12 |
| TRPA | Tryptophan synthase subunit alpha | -2.24 | -0.64 | -0.02 |
| FUMA | Fumarate hydratase class I | -2.23 | -1.65 | -0.13 |
| NDK | Nucleoside diphosphate kinase | -2.2 | -2.04 | -0.31 |
| MASY | Malate synthase A | -2.19 | -2.96 | -0.16 |
| LSRK | Autoinducer-2 kinase | -2.17 | -3.45 | -0.40 |
| TNAA | Tryptophanase | -2.16 | -2.21 | -0.17 |
| DCTA | Aerobic C4-dicarboxylate transport protein | -2.15 | -1.35 | 0.01 |
| CISY | Citrate synthase | -2.11 | -1.98 | -0.08 |
| ACEA | Isocitrate lyase | -2.07 | -2.73 | -0.21 |
| PAAG | 1,2-epoxyphenylacetyl-CoA isomerase | -1.91 | ND | ND |
| ALR2 | Alanine racemase 2 | -1.88 | -1.23 | 0.06 |
| GLPF | Glycerol uptake facilitator protein | -1.88 | -1.64 | 0.01 |

| | | | | |
|---|---|---|---|---|
| ACNB | Aconitate hydratase | -1.87 | -1.63 | -0.06 |
| GLTI | Glutamate/aspartate ABC transporter periplasmic binding | -1.87 | -1.42 | -0.11 |
| YQEF | Putative acyltransferase | -1.82 | -1.55 | -0.14 |
| PAAZ | Oxepin-CoA hydrolase | -1.81 | ND | ND |
| OPPA | Oligopeptide ABC transporter periplasmic binding protein | -1.74 | -1.64 | -0.14 |
| LCFA | Long-chain-fatty-acid--CoA ligase | -1.73 | -2.63 | -0.05 |
| NAPA | Periplasmic nitrate reductase subunit | -1.73 | -0.31 | 0.31 |
| RPPH | RNA pyrophosphohydrolase | -1.70 | ND | ND |
| ODO2 | Succinyltransferase | -1.70 | -1.47 | -0.04 |
| ENTC | Isochorismate synthase | -1.61 | ND | 1.11 |
| XYLA | Xylose isomerase | -1.60 | -0.21 | -0.09 |
| GLPK | Glycerol kinase | -1.58 | -1.39 | -0.06 |
| GLNQ | L-glutamine ABC transporter ATP binding subunit | -1.58 | -1.41 | -0.06 |
| YEGW | Putative DNA-binding transcriptional regulator | -1.57 | -0.75 | 0.05 |
| FADL | Long-chain fatty acid transport | -1.56 | -1.39 | 0.05 |
| HISP | Lysine/arginine/ornithine ABC transporter | -1.52 | -2.74 | -0.14 |
| YTFJ | Unknown function | -1.46 | -0.87 | -0.18 |
| PAAH | 3-hydroxyadipyl-CoA dehydrogenase | -1.46 | ND | ND |
| YEDF | Putative sulfurtransferase | -1.43 | -0.91 | -0.21 |
| FLIC | Flagellar filament structural protein | -1.40 | -0.60 | -0.08 |
| DPPB | Dipeptide ABC transporter membrane subunit | -1.33 | ND | ND |
| GLPT | Sn-glycerol 3-phosphate:phosphate antiporter | -1.32 | -0.88 | 0.04 |
| RIBB | 3,4-dihydroxy-2-butanone-4-phosphate synthase | -1.30 | -0.65 | -0.20 |
| ASNA | Asparagine synthetase A | -1.28 | -2.35 | 0.06 |
| YNJH | Unknown function | -1.24 | -1.57 | -0.18 |
| MALM | Maltose regulon periplasmic protein | -1.21 | -1.11 | -0.11 |
| GLPQ | Glycerophosphoryl diester phosphodiesterase | -1.17 | -0.67 | -0.06 |
| MTFA | Mlc titration factor | -1.17 | -1.02 | -0.18 |
| RIHC | Ribonucleoside hydrolase | -1.16 | -1.11 | -0.36 |
| FLIY | L-cystine-binding protein | -1.16 | -0.76 | -0.18 |
| GSIB | Glutathione ABC transporter periplasmic binding protein | -1.16 | -0.21 | ND |
| OPPF | Oligopeptide transport ATP-binding protein | -1.15 | -0.51 | -0.06 |
| RBSB | Ribose ABC transporter periplasmic binding protein | -1.15 | -1.07 | -0.13 |
| CHEB | Vhemotaxis protein | -1.07 | -0.93 | -0.03 |
| YIDQ | Unknown function | -1.07 | -1.54 | 0.32 |
| CSPD | DNA replication inhibitor | -1.06 | -0.95 | 0.09 |
| PSPE | Tthiosulfate sulfurtransferase | -1.04 | -0.50 | -0.25 |
| TAR | Methyl-accepting chemotaxis protein II | -1.00 | -0.72 | -0.11 |
| NUDL | Unknown function | 6.29 | ND | ND |
| SLP | Starvation lipoprotein, involved in acid resistance | 6.16 | 6.01 | -0.22 |
| HDEA | Acid stress chaperone | 6.02 | 5.59 | 0.16 |

| | | | | |
|---|---|---|---|---|
| DCEB | Glutamate decarboxylase B | 5.5 | 6.91 | 0.27 |
| LDCI | Lysine decarboxylase 1 | 4.79 | 5.38 | 0.49 |
| YMGD | Unknown function | 3.7 | 4.95 | -0.04 |
| OSMB | Osmotically-inducible lipoprotein | 2.9 | 4.18 | 0.26 |
| SFGH1 | S-formylglutathione hydrolase | 2.48 | ND | ND |
| YHAM | Putative L-cysteine desulfidase | 2.45 | 1.28 | 0.75 |
| MLTB | Membrane-bound lytic murein transglycosylase B | 2.38 | 0.47 | 0.06 |
| MLIC | Membrane-bound lysozyme inhibitor | 2.27 | 2.73 | 0.09 |
| EPTA | Phosphoethanolamine transferase | 2.27 | 2.20 | 0.20 |
| MSCS | Small conductance mechanosensitive channel | 2.11 | 0.99 | 0.11 |
| CFA | Cyclopropane fatty acyl phospholipid synthase | 2.10 | 0.98 | 0.03 |
| NUPC | Nucleoside: H + symporter | 2.00 | 0.90 | 0.34 |
| HSLJ | Lipoprotein implicated in Novobiocin resistance | 1.91 | 2.28 | 0.31 |
| YECN | Unknown function | 1.83 | ND | ND |
| APBE | FAD:protein FMN transferase | 1.71 | -0.53 | -0.08 |
| DEOC | Deoxyribose-phosphate aldolase | 1.71 | 1.78 | 0.06 |
| CDD | Cytidine/deoxycytidine deaminase | 1.63 | 1.76 | -0.05 |
| TYPH | Thymidine phosphorylase | 1.6 | 1.59 | -0.04 |
| DEOB | Phosphopentomutase | 1.57 | 1.56 | 0.05 |
| HHA | Hemolysin expression-modulating protein | 1.47 | ND | ND |
| SUFD | Fe-S cluster scaffold complex subunit | 1.39 | ND | ND |
| UDP | UDP-glucose | 1.38 | 2.01 | -0.05 |
| USPD | Universal stress protein D | 1.35 | 1.52 | 0.24 |
| G6PI | Lucose-6-phosphate isomerase | 1.29 | 1.56 | -0.02 |
| KPYK1 | Pyruvate kinase I | 1.28 | 1.57 | 0.12 |
| EDD | Phosphogluconate dehydratase | 1.27 | 0.67 | 0.07 |
| MLTA | Membrane-bound lytic murein transglycosylase A | 1.24 | 0.48 | 0.23 |
| OMPX | Outer membrane protein X | 1.24 | 1.19 | 0.26 |
| ODP2 | Dihydrolipoamide acetyltransferase | 1.23 | 1.43 | 0.04 |
| GPMI | 2,3-bisphosphoglyerate-dependent | 1.13 | 1.35 | 0.12 |
| YIAD | Family lipoprotein | 1.09 | 1.06 | 0.17 |
| TESB | Acyl-CoA thioesterase II | 1.04 | 0.71 | -0.05 |
| FRSA | Fermentation-respiration switch protein | 1.04 | 1.34 | 0.08 |
| RNPH | RNA pyrophosphohydrolase | 1.03 | 1.16 | -0.22 |
| GSP | Glutathionylspermidine synthetase | 1.01 | 1.19 | -0.04 |

ND: not detected

Data from Figures 2B and Figure S3C, S3D. Differences with p-value <0.05 and $Log_2$ of fold change >1 were considered significant.

**Table S3. Proteins showing significant differences in the *ΔhflKC* and wild-type strains during growth in TB at 220 rpm anaerobically**

| | | $Log_2$ (Fold change) |
|---|---|---|
| Protein | Function | *ΔhflKC* vs WT |
| Respiratory proteins | | |
| ISPG | Oxidoreductase involved in isoprenoid biosynthesis | -3.53 |
| UBIT | Anaerobic ubiquinone biosynthesis | -1.52 |
| UBIE | Ubiquinone biosynthesis | -1.17 |
| NARG | Oxidoreductase, nitrate reductase | 1.43 |
| AIDB | Putative acyl-CoA dehydrogenase | 1.39 |
| NARH | Oxidoreductase, nitrate reductase | 1.33 |
| Other proteins | | |
| AG43 | Biofilm formation, Antigen 43 | -3.43 |
| FLHD | Flagellar transcriptional activator | -3.06 |
| ASNA | Asparagine synthetase A | -2.09 |
| HISP | Lysine/arginine/ornithine ABC transporter | -1.72 |
| RBSR | DNA-binding transcriptional repressor | -1.63 |
| ISCR | DNA-binding transcriptional activator | -1.39 |
| YDCI | DNA-binding transcriptional dual regulator | -1.32 |
| YDHC | Putative transporter | -1.30 |
| FIEF | Ferrous-iron efflux pump | -1.28 |
| FLIL | Flagellar biosynthesis protein | -1.20 |
| ILVB | Acetolactate synthase | -1.10 |
| FTSB | Cell division protein | -1.08 |
| NARJ | Molybdenum-cofactor-assembly chaperone | 3.45 |
| FIMH | Minor component of type 1 fimbriae | 3.03 |
| DCEB | Glutamate decarboxylase B | 2.74 |
| FIMG | Minor component of type 1 fimbriae | 2.22 |
| YEHZ | Glycine betaine-binding protein | 1.98 |
| YDJN | Uncharacterized protein | 1.97 |
| DCEA | Glutamate decarboxylase A | 1.96 |
| GADC | Glutamate/gamma antiporter | 1.65 |
| ACPH | Acylaminoacyl-peptidase | 1.57 |
| YMGG | Unknown function | 1.51 |
| HDEA | Acid stress chaperone | 1.48 |
| SLP | Starvation lipoprotein, involved in acid resistance | 1.33 |
| INTA | Prophage integrase | 1.26 |
| YRDB | Uncharacterized protein | 1.25 |
| YNBE | Uncharacterized protein | 1.22 |
| HYAF | Hydrogenase-1 operon protein | 1.17 |

| | | |
|---|---|---|
| MDTF | Multidrug efflux pump | 1.06 |
| FIMF | Minor component of type 1 fimbriae | 1.04 |
| IVY | Inhibitor of vertebrate lysozyme | 1.04 |
| PPA | Inorganic pyrophosphatase | 1.04 |
| HDEB | Acid stress chaperone | 1.03 |

Data from Figure 2C. Differences with p-value <0.05 and $Log_2$ of fold change > 1 were considered significant.

**Table S4. Proteins showing significant differences in *ispG* knockdown and wild-type* strains during growth in TB at 220 rpm**

| | | $Log_2$ (Fold change) |
|---|---|---|
| Protein | Function | *ispG* knockdown[a] vs WT* + pgRNA |
| Respiratory proteins | | |
| NARG | Oxidoreductase, nitrate reductase | -7.99 |
| NIRB | Nitrite Reductase | -5.83 |
| NARH | Oxidoreductase, nitrate reductase | -5.25 |
| NIRD | Nitrite reductase | -5.15 |
| LLDD | L-lactate dehydrogenase | -4.62 |
| ISPG | Oxidoreductase involved in isoprenoid biosynthesis | -4.20 |
| YFCG | Disulfide bond oxidoreductase | -2.84 |
| HYBA | Hydrogenase 2 iron-sulfur protein | -2.77 |
| GLPC | Anaerobic glycerol-3-phosphate dehydrogenase subunit C | -2.65 |
| NAPA | Periplasmic nitrate reductase subunit | -2.47 |
| SDHD | Succinate:quinone oxidoreductase, membrane protein | -1.85 |
| GLPB | Anaerobic glycerol-3-phosphate dehydrogenase subunit B | -1.67 |
| GLPA | Anaerobic glycerol-3-phosphate dehydrogenase subunit A | -1.54 |
| PUTA | Oxidoreductase, proline dehydrogenase | -1.52 |
| MQO | Malate:quinone oxidoreductase | -1.36 |
| STHA | Oxidoreductase, NAD(P)+ transhydrogenase | -1.28 |
| MAO2 | Malate dehydrogenase | -1.27 |
| ALDA | Lactaldehyde dehydrogenase A | -1.25 |
| FRDB | Fumarate reductase iron-sulfur protein | -1.21 |
| FRDA | Fumarate reductase flavoprotein subunit | -1.15 |
| ENTA | 2,3-dihydro-2,3-dihydroxybenzoate dehydrogenase | -1.12 |
| SUCC | Succinyl-CoA synthetase subunit β | -1.12 |
| SDHB | Succinate:quinone oxidoreductase | -1.09 |
| SUCD | Succinyl-CoA synthetase subunit α | -1.03 |

| | | |
|---|---|---|
| SDHA | Succinate:quinone oxidoreductase | -0.98 |
| DHSD | Succinate:quinone oxidoreductase | -0.96 |
| YEIQ | Putative Oxidoreductase | -0.95 |
| DADA | D-amino acid dehydrogenase | -0.90 |
| UBIE | Ubiquinone biosynthesis | -0.88 |
| FADE | Acyl-CoA dehydrogenase | -0.79 |
| CYOA | Cytochrome bo(3) ubiquinol oxidase subunit 2 | -0.78 |
| DHG | Quinoprotein glucose dehydrogenase | -0.70 |
| NRDI | Dimanganese-Tyrosyl Radical Cofactor Maintenance Flavodoxin | -0.70 |
| CYOB | Cytochrome bo(3) ubiquinol oxidase subunit 1 | -0.69 |
| ODO1 | Oxidoreductase, 2-oxoglutarate dehydrogenase | -0.67 |
| YKGF | L-lactate dehydrogenase complex protein | 4.12 |
| YKGE | L-lactate dehydrogenase complex protein | 2.59 |
| SRLD | Sorbitol-6-phosphate 2-dehydrogenase | 1.92 |
| ADHE | Fused acetaldehyde-CoA dehydrogenase | 1.60 |
| FHUF | Ferric-siderophore reductase | 1.58 |
| CYDB | Cytochrome bd-I ubiquinol oxidase subunit 2 | 1.41 |
| DHNA | Type II NADH:quinone oxidoreductase | 1.18 |
| CYDA | Cytochrome bd-I ubiquinol oxidase subunit 1 | 1.16 |
| AIDB | Putative acyl-CoA dehydrogenase | 1.12 |
| IDND | L-idonate 5-dehydrogenase | 1.08 |
| YGHA | NADP(+)-dependent aldehyde reductase | 1.04 |
| POXB | Pyruvate oxidase | 1.01 |
| HISX | Histidinol dehydrogenase | 0.97 |
| LDHD | D-lactate dehydrogenase | 0.97 |
| ADHP | Ethanol dehydrogenase | 0.93 |
| YGHU | Oxidoreductase, organic hydroperoxide reductase | 0.88 |
| CURA | NADPH-dependent curcumin reductase | 0.86 |
| NQOR | NAD(P)H:quinone oxidoreductase | 0.79 |
| GLRX2 | Reduced glutaredoxin 2 | 0.76 |
| GATD | Galactitol-1-phosphate 5-dehydrogenase | 0.67 |
| AHPF | Oxidoreductase, alkyl hydroperoxide reductase | 0.62 |
| YDGJ | Putative oxidoreductase | 0.61 |
| CATE | Oxidoreductases, catalase-peroxidase | 0.81 |

Other proteins

| | | |
|---|---|---|
| LLDP | Lactate/Glycolate:H(+) Symporter | -5.42 |
| DMSB | Dimethyl sulfoxide reductase subunit B | -4.62 |
| NIKA | Ni(2(+)) ABC Transporter periplasmic binding protein | -4.21 |
| NIRC | Nitrite transporter | -3.34 |
| NANT | Sialic acid transporter | -3.13 |

| | | |
|---|---|---|
| YNJE | molybdopterin synthase sulfurtransferase | -3.11 |
| YJJI | Unknown function | -3.05 |
| YIGI | Putative thioesterase | -2.95 |
| YJIY | Pyruvate:H(+) symporter | -2.89 |
| KPTA | RNA 2'-Phosphotransferase | -2.71 |
| MHPR | DNA-binding transcriptional activator | -2.57 |
| ASPG2 | L-asparagine 2 | -2.52 |
| IBPB | Small heat shock protein | -2.39 |
| BIOD2 | Dethiobiotin synthetase | -2.11 |
| FIU | Iron transport | -2.01 |
| CSPG | Cold shock protein | -1.92 |
| NAPH | Ferredoxin-type protein | -1.82 |
| GARD | Galactarate dehydratase | -1.78 |
| FHUE | Ferric coprogen outer membrane receptor | -1.76 |
| GARL | 5-keto-4-deoxy-D-glucarate aldolase | -1.75 |
| FADA | 3-ketoacyl-CoA thiolase | -1.74 |
| BHSA | Multiple stress resistance outer membrane | -1.73 |
| GRCA | Autonomous glycyl radical cofactor | -1.69 |
| YCFS | L,D-transpeptidase | -1.68 |
| FEPA | Ferric enterobactin outer membrane transporter | -1.66 |
| NANK | N-acetylmannosamine kinase | -1.63 |
| HYPD | Hydrogenase maturation factor | -1.57 |
| CIRA | Colicin I receptor | -1.57 |
| FADB | Fatty acid oxidation complex subunit alpha | -1.52 |
| PSTS | Phosphate ABC Transporter periplasmic binding protein | -1.51 |
| NANE | Putative N-acetylmannosamine-6-phosphate 2-epimerase | -1.49 |
| NANA | N-acetylneuraminate lyase | -1.42 |
| DGAL | D-galactose/methyl-galactoside ABC transporter | -1.41 |
| YEGD | Unknown function | -1.41 |
| LIVF | High-affinity branched-chain amino acid transport ATP | -1.41 |
| YBDZ | Enterobactin biosynthesis protein | -1.40 |
| FUMA | Fumarate hydratase class I | -1.39 |
| FUMC | Fumarate hydratase class II | -1.37 |
| YEJG | Unknown function | -1.37 |
| CISY | Citrate synthase | -1.36 |
| ENTH | Proofreading thioesterase | -1.33 |
| IBPA | Small heat shock protein | -1.27 |
| YDCI | DNA-binding transcriptional dual regulator | -1.27 |
| ENTF | Enterobactin synthase component F | -1.26 |
| ENTC | Isochorismate synthase | -1.19 |
| NDK | Nucleoside diphosphate kinase | -1.16 |
| EFEB | Heme-containing peroxidase/deferrochelatase | -1.16 |

| | | |
|---|---|---|
| KDPB | Potassium-transporting ATPase | -1.15 |
| YHCH | N-acetylneuraminate anomerase | -1.14 |
| ENTB | Enterobactin synthase component B | -1.07 |
| YBIX | Unknown function | -1.06 |
| FES | Ferric enterobactin esterase | -1.06 |
| UGPB | Transport of sn-glycerol 3-phosphate | -1.05 |
| YFGM | Ancillary SecTEG translocon subunit | -1.04 |
| MGLA | ATP-binding component of a D-galactose | -1.02 |
| YJIA | P-Loop guanosine triphosphatase | -1.02 |
| PPPA | Prepilin Peptidase | -1.01 |
| YDFZ | Putative Selenoprotein | -1.00 |
| FADJ | Unsaturated acyl-CoA hydratase | -1.00 |
| YBJX | Unknown function | -0.96 |
| ACTP | Acetate transpor | -0.95 |
| PHOQ | Sensor histidine kinase | -0.95 |
| ENTE | Enterobactin synthase component E | -0.94 |
| LIVJ | Branched chain amino acid/phenylalanine ABC | -0.93 |
| MGLC | ATP-binding component of a D-galactose | -0.91 |
| PUTP | Sodium/proline transporter | -0.90 |
| PSTB | Phosphate ABC Transporter ATP Binding Subunit | -0.90 |
| CSPA | Cold shock protein | -0.88 |
| DER | 50S Ribosomal Subunit stability factor | -0.87 |
| FEOA | Ferrous iron transport protein A | -0.86 |
| ACSA | Acetyl-coenzyme A synthetase | -0.86 |
| OMPF | Outer membrane porin F | -0.86 |
| CYCA | D-serine/D-alanine/glycine transporter | -0.85 |
| FEOB | Fe 2 (+) transporter | -0.84 |
| SDAC | L-Serine:H(+) symporter | -0.84 |
| ODO2 | Succinyltransferase | -0.84 |
| GAL7 | Galactose-1-phosphate uridylyltransferase | -0.84 |
| SYH | Histidine--tRNA ligase HisS | -0.83 |
| ALR2 | Alanine racemase 2 | -0.82 |
| LIVK | L-phenylalanine ABC transporter periplasmic binding protein | -0.82 |
| UXAC | D-Glucoronate/D-Galacturonate isomerase | -0.82 |
| FADI | Acetyl-CoA acyltransferase | -0.81 |
| PRIC | Primosomal replication Protein N'' | -0.81 |
| TNAA | Tryptophanase | -0.80 |
| FEOC | Ferrous iron transport protein | -0.80 |
| YDDB | Unknown function | -0.77 |
| BAMB | Outer membrane protein assembly factor | -0.76 |
| YFCZ | Unknown function | -0.75 |
| YNCE | PQQ-like domain-containing protein | -0.74 |

| | | |
|---|---|---|
| RIR3 | Ibonucleoside-diphosphate reductase 2 | -0.74 |
| ACNB | Aconitate hydratase | -0.72 |
| DEAD | ATP-dependent RNA helicase | -0.71 |
| YNCD | Pyrroloquinoline quinone TonB-dependent | -0.70 |
| PEPT | Peptidase T | -0.70 |
| EFEO | Ferrous iron transport system | -0.67 |
| YFAZ | Unknown function | -0.66 |
| ACFD | Putative lipoprotein | -0.66 |
| YFGJ | Zinc ribbon domain-containing protein | -0.65 |
| UXAA | D-Altronate dehydratase | -0.64 |
| T1RK | Type I site-specific deoxyribonuclease | -0.63 |
| LDCI | Lysine decarboxylase 1 | 5.60 |
| RFAS | Lipopolysaccharide core biosynthesis protein | 4.30 |
| YKGG | Unknown function | 2.89 |
| RMF | Ribosome modulation factor | 2.57 |
| OTSB | Trehalose-6-phosphate phosphatase | 2.49 |
| YMGD | Unknown function | 2.24 |
| DPS | DNA protection during starvation protein | 1.96 |
| HDEA | Acid stress chaperone | 1.81 |
| ILVG | Acetolactate synthase isozyme 2 | 1.80 |
| PTHA | Glucitol/sorbitol transport | 1.67 |
| IVY | Inhibitor of vertebrate lysozyme | 1.63 |
| BFR | Bacterioferritin | 1.60 |
| YEBV | Unknown function | 1.50 |
| YHBO | Protein/Nucleic acid deglycase 2 | 1.35 |
| GUTM | DnabBinding transcriptional activator | 1.32 |
| HIS1 | ATP phosphoribosyltransferase | 1.29 |
| GNTK | D-Gluconate Kinase, Thermostable | 1.29 |
| GATR | Putative galactitol utilization operon repressor | 1.23 |
| MLIC | Membrane-bound lysozyme inhibitor | 1.23 |
| EVGS | Sensor histidine kinase | 1.22 |
| MALZ | Maltodextrin glucosidase | 1.16 |
| YGAM | Unknown function | 1.15 |
| FIC | Putative adenosine monophosphate | 1.06 |
| SRA | Ribosome-associated protein | 1.06 |
| HDEB | Acid stress chaperone | 1.05 |
| YEAG | Protein kinase | 1.05 |
| YHJG | Unknown function | 1.05 |
| ELAB | Unknow function | 1.02 |
| ALF | Fructose-bisphosphate aldolase class II | 1.01 |
| AMY1 | Periplasmic alpha-amylase | 1.00 |
| RFAJ | Lipopolysaccharide biosynthesis | 1.00 |
| YDIH | Unknown function | 0.99 |

| | | |
|---|---|---|
| HIS4 | Phosphoribosylformimino- isomerase | 0.98 |
| RPOS | Rna polymerases sigma factor | 0.97 |
| YGAU | K(+) binding protein | 0.95 |
| HIS5 | Imidazole glycerol phosphate synthase | 0.95 |
| TALA | Transaldolase A | 0.94 |
| YCAC | Putative hydrolase | 0.89 |
| TKT2 | Transketolase 2 | 0.89 |
| RFAY | Lipopolysaccharide core heptose (II) kinase | 0.89 |
| HIS8 | Histidinol-phosphate aminotransferase | 0.88 |
| RFAI | LPS alpha-1,3-glucosyltransferase | 0.88 |
| PTND | Mannose-specific PTS enzyme | 0.87 |
| HIS7 | Imidazoleglycerol-phosphate dehydratase | 0.86 |
| GATY | Tagatose-1,6-bisphosphate aldolase 2 | 0.85 |
| DHAK | Dihydroxyacetone kinase subunit K | 0.85 |
| YJBJ | Putative stress response protein | 0.82 |
| RBSC | Ribose ABC Transporter membrane subunit | 0.82 |
| CSIE | Stationary phase-Iinducible protein | 0.80 |
| MPAA | Murein tripeptide amidase A | 0.80 |
| SLP | Starvation lipoprotein, involved in acid resistance | 0.79 |
| ALF1 | Fructose-bisphosphate aldolase class I | 0.79 |
| ILVC | Ketol-acid reductoisomerase | 0.78 |
| HIS6 | Imidazole glycerol phosphate synthase | 0.77 |
| PTHB | Glucitol/sorbitol transport | 0.76 |
| PGK | Phosphoglycerate kinase | 0.76 |
| GUTQ | D-arabinose 5-phosphate isomerase | 0.75 |
| YAIA | Unknown function | 0.74 |
| USPD | Universal stress protein D | 0.74 |
| OTSA | Trehalose-6-phosphate synthase | 0.73 |
| TREF | Cytoplasmic trehalase | 0.72 |
| BEPA | Beta-barrel assembly-enhancing protease | 0.72 |
| FABR | Dna-binding transcriptional repressor | 0.71 |
| GSP | Glutathionylspermidine synthetase | 0.70 |
| TREA | Periplasmic trehalase | 0.70 |
| GLK | Glucokinase | 0.69 |
| MDTE | Multidrug efflux pump | 0.69 |
| YGDI | Unknown function | 0.69 |
| PTNAB | Mannose-specific PTS enzyme | 0.69 |
| NPD | NAD-dependent deacetylase | 0.69 |
| G6PI | Lucose-6-phosphate isomerase | 0.68 |
| DHAL | Dihydroxyacetone kinase subunit L | 0.67 |
| MSYB | Acidic protein | 0.66 |
| YFBU | Unknown function | 0.66 |
| DHAM | Dihydroxyacetone kinase subunit M | 0.65 |

| | | |
|---|---|---|
| YCJW | DNA-binding transcriptional repressor | 0.64 |
| YCEF | 7-methyl-GTP pyrophosphatase | 0.64 |
| RBSA | Ribose ABC Transporter ATP binding subunit | 0.64 |
| GSIA | Glutathione ABC Transporter ATP Binding Subunit | 0.63 |
| ECNB | Bacteriolytic entericidin B lipoprotein | 0.63 |
| TAM | Trans-aconitate 2-methyltransferase | 0.63 |
| YBHB | Putative kinase inhibitor | 0.63 |
| AK3 | Aspartate kinase III | 0.63 |
| YJDC | Putative DNA-binding transcriptional regulator | 0.63 |
| PTKC | Galactitol-specific enzyme | 0.62 |
| YGIM | Putative signal transduction protein | 0.62 |
| COBS | Cobalamin 5'-Phosphate synthase | 0.62 |
| YBGS | Unknown function | 0.62 |
| EDD | Phosphogluconate dehydratase | 0.61 |
| LPXC | N-Acetylglucosamine deacetylase | 0.61 |
| YHEV | Duf2387 domain-containing Protein | -0.61 |

* *E. coli* YYdCas9

[a]*E. coli* YYdCas9 + pMI112 induced with 0.2 µM aTC

Data from Figure 4I. Differences with p-value <0.06 and $Log_2$ of fold change > 2 were considered significant.

**Table S5. Proteins showing significant differences in *ΔarcB* and wild-type strains during growth in TB at 220 rpm**

| | | $Log_2$ (Fold change) |
|---|---|---|
| Protein | Function | *ΔarcB* vs WT |
| Respiratory proteins | | |
| NIRB | Nitrite reductase (NADH) large subunit | -5.72 |
| NAPA | Periplasmic nitrate reductase subunit | -4.21 |
| QUEG | Epoxyqueuosine reductase | -3.68 |
| DMSB | Dimethyl sulfoxide reductase subunit B | -3.42 |
| CYDA | Cytochrome bd-I ubiquinol oxidase subunit 1 | -2.70 |
| CYDB | Cytochrome bd-I ubiquinol oxidase subunit 2 | -2.63 |
| UBIT | Anaerobic ubiquinone biosynthesis | -2.56 |
| YAHK | Aldehyde reductase, NADPH-dependent | 2.02 |
| HISX | Histidinol dehydrogenase | 2.03 |
| PUTA | Oxidoreductase, proline to glutamate | 2.07 |
| GPR | L-glyceraldehyde 3-phosphate reductase | 2.08 |
| FADJ | Unsaturated acyl-CoA hydratase | 2.08 |
| NUOH | NADH:quinone oxidoreductase subunit H | 2.10 |
| ACDH | Acetaldehyde dehydrogenase | 2.22 |

| | | |
|---|---|---|
| NUOM | NADH:quinone oxidoreductase subunit M | 2.32 |
| YCAK | Putative oxidoreductase | 2.37 |
| YBIC | Hydroxycarboxylate dehydrogenase B | 2.45 |
| HCAD | Putative 3-phenylpropionate | 2.51 |
| IDH | Oxidoreductase, isocitrate dehydrogenase | 2.54 |
| MQO | Malate:quinone oxidoreductase | 2.55 |
| POXB | Pyruvate oxidase | 2.61 |
| MDH | Malate dehydrogenase | 2.66 |
| CURA | NADPH-dependent curcumin reductase | 2.89 |
| ADHE | Fused acetaldehyde-CoA dehydrogenase | 2.93 |
| SDHB | Succinate:quinone oxidoreductase | 3.03 |
| SDHA | Succinate:quinone oxidoreductase | 3.07 |
| DHSD | Succinate:quinone oxidoreductase | 3.15 |
| ODO2 | Dihydrolipoyltranssuccinylase | 3.20 |
| SUCD | Succinyl-CoA synthetase subunit α | 3.28 |
| ALDA | Aldehyde dehydrogenase A | 3.31 |
| SUCC | Succinyl-CoA synthetase subunit β | 3.34 |
| DADA | D-amino acid dehydrogenase | 3.52 |
| IDND | L-idonate 5-dehydrogenase | 3.82 |
| ABDH | Ethanol dehydrogenase/alcohol dehydrogenase | 3.82 |
| LLDD | L-lactate dehydrogenase | 4.04 |
| FADE | Acyl-CoA dehydrogenase | 4.12 |
| GABD | Succinate-semialdehyde dehydrogenase | 4.24 |
| YKGE | L-lactate dehydrogenase complex protein | 4.40 |
| LHGO | L-2-hydroxyglutarate dehydrogenase | 4.53 |
| SRLD | Sorbitol-6-phosphate 2-dehydrogenase | 5.14 |
| SODC | Superoxide dismutase (Cu-Zn) | 5.15 |
| YKGF | L-lactate dehydrogenase complex protein | 5.35 |
| ASTD | Aldehyde dehydrogenase | 6.07 |
| CSID | Oxidoreductase, glutarate hydroxylase | 8.50 |

Other proteins

| | | |
|---|---|---|
| YEER | CP4-44 prophage | -5.89 |
| NIRD | Nitrite reductase | -5.17 |
| TDCB | Catabolic threonine dehydratase | -5.00 |
| NIRC | Nitrite transporter | -4.99 |
| YFDC | Unknown function | -4.28 |
| AG43 | Biofilm formation, Antigen 43 | -4.20 |
| ADEP | Adenine:H (+) symporter | -3.93 |
| PLAP | Putrescine:H (+) symporter | -3.41 |
| YGIQ | Unknown function | -3.33 |
| YGHG | Lipoprotein | -3.31 |

| | | |
|---|---|---|
| MEND | Menaquinone pathway | -3.22 |
| ACFD | Putative lipoprotein | -2.86 |
| CODB | Cytosine transporter | -2.75 |
| RECN | DNA repair protein | -2.64 |
| YDCP | 23S rRNA 5-hydroxycytidine C2501 synthase | -2.63 |
| QSEC | Sensor histidine kinase | -2.59 |
| FEOB | Fe 2 (+) transporter | -2.45 |
| CSPB | Cold shock-like protein | -2.43 |
| FTNB | Putative ferritin-like protein | -2.43 |
| CSPA | Cold shock protein | -2.42 |
| PYRE | Orotate phosphoribosyltransferase | -2.41 |
| YBFE | Unknown function | -2.39 |
| FIS | DNA-binding transcriptional dual regulator | -2.37 |
| TADA | tRNA adenosine (34) deaminase | -2.35 |
| YDIY | Unknown function | -2.33 |
| YJJI | Unknown function | -2.24 |
| BCSE | c-di-GMP-binding protein | -2.24 |
| EPMA | Enlongation factor P | -2.23 |
| GHXP | Guanine/hypoxanthine transporter | -2.19 |
| DEAD | ATP-dependent RNA helicase | -2.16 |
| HYPD | Hydrogenase maturation factor | -2.12 |
| FADL | Long-chain fatty acid transport protein | -2.09 |
| RLMG | rRNA base methyltransferase | -2.06 |
| RFAY | Lipopolysaccharide core heptose (II) kinase | -2.06 |
| KDPB | Potassium-transporting ATPase | -2.06 |
| RAPA | RNA polymerase-binding ATPase | -2.06 |
| RHLE | ATP-dependent RNA helicase | -2.05 |
| FEOA | Ferrous iron transport protein A | -2.03 |
| ANSP | L-asparagine transporter | -2.02 |
| LDCI | Lysine decarboxylase 1 | 8.63 |
| ASTC | Succinylornithine aminotransferase | 7.46 |
| YEAG | Protein kinase | 5.46 |
| ACTP | Acetate transpor | 5.41 |
| RPOS | RNA polymerase sigma factor | 5.37 |
| RMF | Ribosome modulation factor | 5.30 |
| POTF | Putrescine ABC transporter periplasmic binding protein | 5.11 |
| FADB | Fatty acid oxidation complex subunit alpha | 5.03 |
| GABT | 4-aminobutyrate aminotransferase | 4.83 |
| PSIF | Phosphate starvation-inducible protein | 4.83 |
| ACSA | Acetyl-coenzyme A synthetase | 4.77 |
| LLDR | L-lactate dehydrogenase operon regulator | 4.74 |
| PTHA | Glucitol/sorbitol transport | 4.56 |
| YGAM | Unknown function | 4.56 |

| | | |
|---|---|---|
| YEBF | Unknown function | 4.49 |
| UGPC | sn-glycerol 3-phosphate ABC transporter | 4.43 |
| PTHC | Glucitol/sorbitol transport | 4.22 |
| PTHB | Glucitol/sorbitol transport | 4.13 |
| YKGG | Unknown function | 4.12 |
| PUUA | Glutamate-putrescine ligase | 4.09 |
| METE | Novo methionine biosynthesis | 4.06 |
| YDCS | Putative ABC transporter periplasmic binding | 4.05 |
| YDIH | Unknown function | 4.03 |
| YJHG | D-xylonate dehydratase | 3.97 |
| FADA | 3-ketoacyl-CoA thiolase | 3.93 |
| ASTA | Arginine N-succinyltransferase | 3.93 |
| CISY | Citrate synthase | 3.83 |
| FUCA | L-fuculose-phosphate aldolase | 3.81 |
| DPPF | Dipeptide ABC transporter ATP binding subunit | 3.81 |
| AMY2 | Alpha-amylase | 3.69 |
| YDEN | Unknown function | 3.63 |
| OTC1 | Ornithine carbamoyltransferase | 3.61 |
| PAAK | Phenylacetate-CoA ligase | 3.58 |
| UGPB | Transport of sn-glycerol 3-phosphate | 3.57 |
| PAAJ | β-ketoadipyl-CoA thiolase | 3.56 |
| MGLA | ATP-binding component of a D-galactose | 3.53 |
| GLCC | DNA-binding transcriptional dual regulator | 3.49 |
| BFR | Bacterioferritin | 3.49 |
| LSRF | 3-hydroxy-2,4-pentadione 5-phosphate thiolase | 3.38 |
| BLUF | Temperature-regulated antirepressor | 3.35 |
| MGLC | ATP-binding component of a D-galactose | 3.20 |
| IDNK | D-gluconate kinase, thermosensitive | 3.19 |
| YFFR | Unknown function | 3.17 |
| AES | Acetylesterase | 3.17 |
| YDEI | BOF family protein | 3.12 |
| FLGJ | Putative peptidoglycan hydrolase | 3.10 |
| MHPC | 2-hydroxy-6-ketonona-2,4-dienedioate hydrolase | 3.07 |
| GALS | DNA-binding transcriptional dual regulator | 3.06 |
| ASTE | Succinylglutamate desuccinylase | 2.99 |
| YHJG | Unknown function | 2.98 |
| ALR2 | Alanine racemase 2 | 2.97 |
| AGAL | Alpha-galactosidase | 2.96 |
| ARGT | Lysine, arginine and ornithine transport | 2.90 |
| FUMC | Fumarate hydratase class II | 2.88 |
| YAHO | Unknown function | 2.87 |
| DGAL | D-galactose/methyl-galactoside ABC transporter | 2.86 |

| | | |
|---|---|---|
| KBAZ | Putative tagatose-1,6-bisphosphate aldolase 1 chaperone | 2.86 |
| PHSM | Maltodextrin phosphorylase | 2.81 |
| FUCK | L-fuculokinase | 2.75 |
| XYLF | Xylose ABC transporter periplasmic binding protein | 2.73 |
| ALF1 | Fructose-bisphosphate aldolase class I | 2.72 |
| DPPA | Dipeptide ABC transporter periplasmic binding protein | 2.71 |
| FRLB | Fructoselysine 6-phosphate deglycase | 2.70 |
| TREA | Periplasmic trehalase | 2.70 |
| DPPC | Dipeptide ABC transporter membrane subunit | 2.68 |
| YBGS | Unknown function | 2.67 |
| OTSA | Trehalose-6-phosphate synthase | 2.66 |
| SRA | Ribosome-associated protein | 2.64 |
| CSPD | DNA replication inhibitor | 2.64 |
| YIGI | Putative thioesterase | 2.61 |
| FUMA | Fumarate hydratase class I | 2.61 |
| YNIA | Putative kinase | 2.59 |
| YEBV | Unknown function | 2.58 |
| MTFA | Mlc titration factor | 2.55 |
| OSMY | Periplasmic chaperone | 2.53 |
| PGPC | Phosphatidylglycerophosphatase C | 2.50 |
| TALA | Transaldolase A | 2.49 |
| YTFQ | Galactofuranose ABC transporter binding protein | 2.48 |
| PHOH | ATP-binding protein PhoH | 2.48 |
| YJHH | Putative 2-dehydro-3-deoxy-D-pentonate aldolase | 2.44 |
| YDCI | DNA-binding transcriptional dual regulator | 2.43 |
| PTKA | PTS system galactitol-specific EIIA component | 2.42 |
| YQHA | Unknown function | 2.40 |
| HISQ | Histidine transport permease | 2.38 |
| ELAB | Unknow function | 2.36 |
| CLSC | Cardiolipin synthase C | 2.36 |
| YBAY | Unknown function | 2.36 |
| GCSP | Glycine decarboxylase | 2.34 |
| PAAB | Phenylacetyl-CoA 1,2-epoxidase subunit B | 2.33 |
| RBBA | Ribosome-associated ATPase | 2.32 |
| ARAF | Arabinose ABC transporter periplasmic | 2.32 |
| MALQ | 4-alpha-glucanotransferase | 2.30 |
| PTKB | PTS system galactitol-specific EIIB component | 2.30 |
| CUTA | Copper binding protein | 2.29 |
| TRG | Methyl-accepting chemotaxis protein | 2.28 |
| IDNO | 5-keto-D-gluconate 5-reductase | 2.27 |
| ECNB | Bacteriolytic entericidin B lipoprotein | 2.27 |
| YEGP | Unknown function | 2.26 |

| | | |
|---|---|---|
| TAM | Trans-aconitate 2-methyltransferase | 2.26 |
| ACNA | Aconitate hydratase 1 | 2.23 |
| CFA | Cyclopropane fatty acyl phospholipid synthase | 2.22 |
| FADI | Acetyl-CoA acyltransferase | 2.22 |
| TKT2 | Transketolase 2 | 2.22 |
| PTTBC | Rehalose phosphotransferase | 2.21 |
| GLGB | 1,4-alpha-glucan branching enzyme | 2.18 |
| TNAA | Tryptophanase | 2.18 |
| OPPB | Murein tripeptide ABC transporter | 2.18 |
| GARL | 5-keto-4-deoxy-D-glucarate aldolase | 2.17 |
| MALE | Maltose binding protein | 2.13 |
| YEGR | Unknown function | 2.12 |
| YODC | Unknown function | 2.10 |
| FLGM | Anti-sigma factor for FliA | 2.06 |
| TAR | Methyl-accepting chemotaxis protein | 2.03 |
| DPPD | Dipeptide ABC transporter ATP binding subunit | 2.03 |
| GATY | Tagatose-1,6-bisphosphate aldolase 2 | 2.02 |
| OMPF | Outer membrane porin F | 2.02 |
| TREC | Trehalose-6-phosphate hydrolase | 2.01 |

Data from Figure 5C. Differences with p-value <0.05 and $Log_2$ of fold change > 2 were considered significant.

**Table S6. Proteins showing significant differences in *ΔhflKC ΔarcB* and *ΔarcB* strains during growth in TB at 220 rpm**

| | | $Log_2$ (Fold change) |
|---|---|---|
| Protein | Function | *ΔhflKC ΔarcB* vs *ΔarcB* |
| Respiratory proteins | | |
| YAHA | Aldehyde reductase, Nadph-dependent | -2.62 |
| UBIT | Anaerobic ubiquinone biosynthesis | -2.57 |
| UXAB | Tagaturonate reductase | -2.53 |
| ISPG | Oxidoreductase involved in isoprenoid biosynthesis | -2.33 |
| UBIE | Ubiquinone biosynthesis | -1.64 |
| PUTA | Oxidoreductase, proline dehydrogenase | -1.08 |
| CYSI | Sulfite reductase, hemoprotein subunit | 1.13 |
| AEGA | Putative oxidoreductase | 1.13 |
| RCLA | Cupric reductase | 1.14 |
| DKGA | Methylglyoxal reductase | 1.16 |
| GLRX2 | Reduced glutaredoxin 2 | 1.23 |
| HDHA | 7-Alpha-hydroxysteroid dehydrogenase | 1.30 |
| YAHK | Aldehyde reductase | 1.43 |

| | | |
|---|---|---|
| YQJG | Glutathionyl-hydroquinone reductase | 1.46 |
| ABDH | Ethanol dehydrogenase/alcohol dehydrogenase | 1.48 |
| ALDB | Aldehyde dehydrogenase B | 1.67 |
| OSMC | Osmotically inducible peroxiredoxin | 1.81 |
| POXB | Pyruvate oxidase | 1.84 |
| CURA | NADPH-dependent curcumin reductase | 1.87 |
| CATE | Oxidoreductases, catalase-peroxidase | 2.03 |
| NQOR | NAD(P)H:quinone oxidoreductase | 2.11 |
| ADHP | Ethanol dehydrogenase | 2.89 |
| YGHA | NADP(+)-dependent aldehyde reductase | 3.98 |
| YBDR | Putative Zn(2(+))-dependent alcohol dehydrogenase | 4.95 |
| FIXB | Putative electron transfer flavoprotein | 5.12 |

Other proteins

| | | |
|---|---|---|
| PLAP | Putrescine:H (+) symporter | -5.04 |
| YHCN | Unknown function | -3.98 |
| THIC | Phosphomethylpyrimidine synthase | -3.47 |
| PFLD | Putative formate acetyltransferase 2 | -2.82 |
| CODB | Cytosine transporter | -2.25 |
| FLGA | Flagellar basal body P-ring formation protein | -2.16 |
| LDCI | Lysine decarboxylase 1 | -1.99 |
| FLHD | Flagellar transcriptional activator | -1.86 |
| CRFC | Regulator of diguanylate cyclase | -1.85 |
| FLHC | DNA-binding transcriptional dual regulator | -1.83 |
| FLGI | Flagellar P-ring protein | -1.75 |
| FHUE | Ferric coprogen outer membrane receptor | -1.72 |
| FIU | Iron transport | -1.70 |
| AER | Aerotaxis receptor | -1.65 |
| CIRA | Colicin I receptor | -1.55 |
| PTOCB | Maltose phosphotransferase | -1.54 |
| ADEC | Adenine deaminase | -1.52 |
| FLGL | Flagellar hook-filament junction protein 2 | -1.51 |
| FLHE | Flagellar protein | -1.49 |
| FLIK | Flagellar hook-length control protein | -1.47 |
| CLSC | Cardiolipin synthase C | -1.42 |
| YDIY | Unknown function | -1.40 |
| MOTB | Flagellar rotation | -1.33 |
| YECR | Unknown function | -1.33 |
| PTTBC | Rehalose phosphotransferase | -1.32 |
| NANA | N-acetylneuraminate lyase | -1.28 |
| SDAC | L-Serine:H (+) symporter | -1.25 |

| | | |
|---|---|---|
| YBIJ | Unknown function | -1.24 |
| FLGE | Flagellar hook protein | -1.23 |
| FLIJ | Flagellar biosynthesis protein | -1.22 |
| OMPF | Outer membrane porin F | -1.21 |
| YDDA | ABC Transporter family protein | -1.20 |
| TNAA | Tryptophanase | -1.20 |
| FLIM | Flagellar motor switch protein | -1.19 |
| YJIY | Pyruvate:H (+) Symporter | -1.17 |
| FLIH | Flagellar biosynthesis protein | -1.17 |
| YBJX | Unknown function | -1.16 |
| FLGD | Flagellar biosynthesis, Initiation of hook assembly | -1.14 |
| FLGF | Flagellar basal-body rod protein | -1.13 |
| FLIS | Flagellar biosynthesis protein | -1.13 |
| TRG | Methyl-accepting chemotaxis protein | -1.13 |
| FLIO | Flagellar biosynthesis protein | -1.13 |
| TREC | Trehalose-6-phosphate hydrolase | -1.12 |
| TSR | Methyl-accepting chemotaxis protein | -1.11 |
| PSUG | Pseudouridine-5'-phosphate glycosidase | -1.11 |
| CSTA | Pyruvate transporter | -1.10 |
| YHJE | Inner membrane metabolite transport protein | -1.10 |
| FLGJ | Putative peptidoglycan hydrolase | -1.10 |
| TRUC | tRNA pseudouridine (65) synthase | -1.09 |
| FLIC | Flagellar filament structural protein | -1.07 |
| PUTP | Sodium/proline transporter | -1.05 |
| DCTA | Aerobic C4-dicarboxylate transport protein | -1.04 |
| BGLR | Beta-glucuronidase | -1.03 |
| PQQL | Periplasmic metalloprotease | -1.02 |
| GLPT | Sn-glycerol 3-phosphate:phosphate antiporter | -1.00 |
| GARD | Galactarate dehydratase | -1.00 |
| FSAB | Fructose-6-phosphate aldolase 2 | -1.00 |
| DGAL | D-galactose/methyl-galactoside ABC transporter | -1.00 |
| GM4D | Mannose 4,6-Dehydratase | 7.33 |
| YBHP | Unknown function | 4.96 |
| YHBO | Protein/Nucleic acid deglycase 2 | 4.51 |
| YEGS | Lipid Kinase | 4.02 |
| UDG | Glucose 6-dehydrogenase | 3.88 |
| OTSB | Trehalose-6-phosphate phosphatase | 3.24 |
| YGDI | Unknown function | 3.01 |
| YBGS | Unknown function | 2.93 |
| PHNB | Unknown function | 2.88 |
| YPFG | Unknown function | 2.85 |
| OSMY | Periplasmic chaperone | 2.65 |
| IDI | Isopentenyl-diphosphate delta-isomerase | 2.55 |

| | | |
|---|---|---|
| MDTE | Multidrug efflux pump | 2.42 |
| YCAC | Putative hydrolase | 2.34 |
| MANB | Phosphomannomutase | 2.30 |
| SRA | Ribosome-associated Protein | 2.29 |
| YCIF | Unknown function | 2.26 |
| YEAG | Protein kinase | 2.26 |
| YGAM | Unknown function | 2.26 |
| OTSA | Trehalose-6-phosphate synthase | 2.24 |
| YGAC | Unknown function | 2.24 |
| YFDC | Unknown function | 2.22 |
| TKT2 | Transketolase 2 | 2.22 |
| DOSC | Diguanylate cyclase | 2.19 |
| IVY | Inhibitor of vertebrate lysozyme | 2.17 |
| DCEA | Glutamate decarboxylase A | 2.13 |
| YDEI | Unknown function | 2.05 |
| TALA | Transaldolase A | 2.03 |
| ELAB | Unknow function | 1.99 |
| YEGP | Unknown function | 1.96 |
| CBPA | Curved DNA-binding protein | 1.88 |
| BFR | Bacterioferritin | 1.87 |
| YJBJ | Putative stress response protein | 1.87 |
| YCCJ | Unknown function | 1.85 |
| YAIA | Unknown function | 1.84 |
| ALF1 | Fructose-bisphosphate aldolase class 1 | 1.83 |
| PHNO | Aminoalkylphosphonate N-acetyltransferase | 1.83 |
| YEHZ | Glycine betaine-binding protein | 1.79 |
| HDEB | Acid stress chaperone | 1.78 |
| MSYB | Acidic protein | 1.76 |
| ECNB | Bacteriolytic entericidin B lipoprotein | 1.76 |
| GCS2 | Carboxylate-amine ligase | 1.76 |
| WZC | Protein-tyrosine kinase | 1.74 |
| YDHS | Unknown function | 1.74 |
| PSIF | Phosphate starvation-inducible protein | 1.73 |
| DPS | DNA protection during starvation protein | 1.68 |
| GADC | Glutamate/gamma antiporter | 1.66 |
| OSME | Osmotically-inducible lipoprotein | 1.66 |
| YBIO | Moderate conductance mechanosensitive channel | 1.62 |
| YAHO | Unknown function | 1.62 |
| YGIW | Cellular response to hydrogen peroxide | 1.60 |
| YBAY | Unknown function | 1.59 |
| MLIC | Membrane-bound lysozyme inhibitor | 1.58 |
| YEBF | Unknown function | 1.57 |
| FIC | Putative adenosine monophosphate | 1.56 |

| | | |
|---|---|---|
| TAM | Trans-aconitate 2-methyltransferase | 1.56 |
| HCHA | Protein/Nucleic acid deglycase 1 | 1.56 |
| DCEB | Glutamate decarboxylase B | 1.55 |
| CSIR | DNA-binding transcriptional repressored Protein | 1.53 |
| YNHG | D-Transpeptidase | 1.48 |
| TREA | Periplasmic trehalase | 1.43 |
| RMF | Ribosome modulation factor | 1.43 |
| INAA | Putative lipopolysaccharide kinase | 1.42 |
| YOBB | Unknown function | 1.36 |
| YHHA | Unknown function | 1.36 |
| YBJP | Unknown function | 1.34 |
| SLP | Starvation lipoprotein, involved in acid resistance | 1.32 |
| YMGG | Unknown function | 1.27 |
| YHJG | Unknown function | 1.27 |
| NHOA | Arylamine N-acetyltransferase | 1.25 |
| YQJE | Unknown function | 1.24 |
| YDIH | Unknown function | 1.23 |
| YODC | Unknown function | 1.22 |
| YQJD | Unknown function | 1.22 |
| YNIA | Putative kinase | 1.19 |
| AG43 | Biofilm formation, Antigen 43 | 1.17 |
| YJDJ | Putative N-acetyltransferase | 1.13 |
| YGAU | K (+) binding protein | 1.12 |
| YDCS | Putative ABC transporter periplasmic binding | 1.11 |
| YOAC | Unknown function | 1.09 |
| CSIE | Stationary phase-inducible protein | 1.09 |
| CYSP | Thiosulfate/Sulfate AbBC Transporter Periplasmic Binding Protein | 1.08 |
| PFKB | 6-Phosphofructokinase 2 | 1.05 |
| GABT | 4-aminobutyrate aminotransferase | 1.03 |

Data from Figure 5E. Differences with p-value <0.05 and $Log_2$ of fold change > 1 were considered significant.

**Table S7. Transcript level of *ispG* and ssrA in *ΔhflKC* and wild-type strains during growth in LB at 220 rpm**

| RNA samples[a] | Target gene | Mean of the Cq value[b] |
|---|---|---|
| WT_A | *ispG* | 22.58 |
| WT_B | | 24.08 |
| WT_C | | 22.33 |
| *ΔhflKC*_A | | 22.61 |
| *ΔhflKC*_B | | 23.81 |
| *ΔhflKC*_C | | 23.30 |
| WT_A | *ssrA* | 12.78 |
| WT_B | | 11.98 |
| WT_C | | 13.55 |
| *ΔhflKC*_A | | 13.28 |
| *ΔhflKC*_B | | 12.33 |
| *ΔhflKC*_C | | 12.72 |
| NRT_WT_A | *ispG* | 33.09 |
| NRT_WT_B | | 37.70 |
| NRT_WT_C | | 37.68 |
| NRT_*ΔhflKC*_A | | 34.18 |
| NRT_*ΔhflKC*_B | | 30.95 |
| NRT_*ΔhflKC*_C | | 36.07 |

[a]Letter A, B, and C represent independent RNA samples.

[b]The relative mRNA level of *ispG* is quantified as the Cq. Data represent four independent replicates. Threshold of the Cq value: 30 cycles

NRT: No Reverse Transcriptase
